# Revisiting chromatin packaging in mouse sperm

**DOI:** 10.1101/2022.12.26.521943

**Authors:** Qiangzong Yin, Chih-Hsiang Yang, Olga S. Strelkova, Jingyi Wu, Yu Sun, Sneha Gopalan, Liyan Yang, Job Dekker, Thomas G. Fazzio, Xin Zhiguo Li, Johan Gibcus, Oliver J. Rando

## Abstract

Mammalian sperm exhibit an unusual and heavily-compacted genomic packaging state. In addition to its role in organizing the compact and hydrodynamic sperm head, it has been proposed that sperm chromatin architecture helps to program gene expression in the early embryo. Scores of genome-wide surveys in sperm have reported patterns of chromatin accessibility, histone localization, histone modification, and chromosome folding. Here, we revisit these studies in light of recent reports that sperm obtained from the mouse epididymis are contaminated with low levels of cell-free chromatin. In the absence of proper sperm lysis we readily recapitulate multiple prominent genome-wide surveys of sperm chromatin, suggesting that these profiles primarily reflect contaminating cell-free chromatin. Removal of cell-free DNA, along with appropriate lysis conditions, are required to reveal a sperm chromatin state distinct from most previous reports. Using ATAC-Seq to explore relatively accessible genomic loci, we identify a landscape of open loci associated with early development and transcriptional control. Histone modification and chromosome folding studies also strongly support the hypothesis that prior studies suffer from contamination, but technical challenges associated with reliably preserving the architecture of the compacted sperm head prevent us from confidently assaying true localization patterns for these epigenetic marks. Together, our studies strongly argue that our knowledge of mammalian chromosome packaging remains largely incomplete, and motivate future efforts to more accurately characterize genome organization in mature sperm.

## INTRODUCTION

The packaging of an organism’s genome into the limiting confines of a typical nucleus requires multiple levels of organization, allowing relatively free access to gene regulatory elements in the face of extensive physical compaction. In eukaryotes, the key principles of chromatin organization have been elucidated over the past half-century: a repeating nucleoprotein complex known as the nucleosome serves to compact ∼150 bp of DNA, resulting in the famous “beads on a string” polymer that is then further organized via loop extrusion and phase separation into larger scale structures such as topologically-associating domains (TADs) and genome “compartments” (Dekker and Misteli 2015; Friedman and Rando 2015; Klemm et al. 2019).

Although we have a relatively mature understanding of chromatin organization in typical somatic cell types, genomic organization in the highly compact sperm nucleus remains less well understood. At the most basic level, it has been known for decades that the vast majority of histones are removed from the genome during the process of spermatogenesis in mammals, to be replaced with small highly basic peptides known as protamines (Bellve et al. 1975; Calvin 1976; Balhorn 1982). Intriguingly, a small subset of nucleosomes are retained in mature sperm – between ∼2% and ∼15%, depending on the mammal (Gaucher et al. 2010; Moritz and Hammoud 2022). A number of studies have mapped the locations of retained nucleosomes, with the majority of reports in humans and mice finding nucleosomes retained at CpG island regulatory elements associated with key developmental genes (Arpanahi et al. 2009; Hammoud et al. 2009; Brykczynska et al. 2010; Erkek et al. 2013; Johnson et al. 2016; Jung et al. 2017; Jung et al. 2019), although a handful of studies instead find histones retained primarily in long stretches over “gene deserts” of low gene density (Carone et al. 2014; Yamaguchi et al. 2018). It appears at least one explanation for this discrepancy may be the extent of nuclease digestion used in mapping nucleosomes in sperm, although many other technical details – including fixation and “pre-swelling” steps prior to chromatin preparation – also differ between studies. Other features of sperm genome compaction that have been investigated in detail include covalent modifications of histones (Hammoud et al. 2009; Brykczynska et al. 2010; Erkek et al. 2013; Jung et al. 2017; Jung et al. 2019; Lismer et al. 2020; Yoshida et al. 2020; Lismer et al. 2021; Bedi et al. 2022), localization of various non-histone proteins (Jung et al. 2017; Jung et al. 2019), and 3-dimensional folding of the sperm genome (Battulin et al. 2015; Jung et al. 2017; Ke et al. 2017; Alavattam et al. 2019; Jung et al. 2019; Vara et al. 2019; Wang et al. 2019b; Gou et al. 2020). In addition to the inherent interest in understanding how the genome folds under conditions of extreme compaction, it has been widely suggested that the organization of the sperm genome – the locations and modifications of retained histones, most notably – may also play a role in programming early gene expression in the preimplantation embryo and may thereby influence later phenotypes in offspring (Siklenka et al. 2015; Lesch et al. 2019).

In the mouse system, sperm are typically collected from the distal, or cauda, region of the epididymis, an epithelial tube where sperm mature for ∼10 days following testicular spermatogenesis. Interestingly, several groups have recently reported the presence of abundant cell-free chromatin in the proximal, or caput, epididymis (Galan et al. 2021; Chen et al. 2022). Although both of these studies focused on this contamination in the caput epididymis, where it results in ∼20-30% contamination of sperm cytosine methylation profiles, there is also a small amount of contaminating chromatin present in cauda sperm preparations – we typically observe ∼2-5% contamination of these sperm preps; the extent of contamination differs somewhat from investigator to investigator. Importantly, ∼2% contamination of cytosine methylation data is comparable to the precision of a typical cytosine methylation survey and is therefore essentially insignificant for most purposes. However, we reasoned that this contamination could be a much greater problem for measurements of sperm chromatin, as 2% genome equivalents of a fully nucleosomal genome would be comparable to the bona fide nucleosomal complement of mouse sperm.

Here, we explored the hypothesis that many or most published studies of mouse sperm chromatin organization reflect contaminating cell free chromatin. Overall, our data strongly imply that current views of the mouse sperm chromatin landscape are flawed, and motivate renewed focus on this important area of chromosome biology and epigenetics.

## RESULTS

### Probing sperm chromatin architecture by ATAC-Seq

In the course of studies focused on dietary effects on sperm chromatin, we carried out ATAC-Seq (<u>A</u>ssay for <u>T</u>ransposase-<u>A</u>ccessible <u>C</u>hromatin by <u>Seq</u>uencing – (Buenrostro et al. 2015)) to identify relatively accessible genomic loci across the sperm genome. Here and through the rest of this manuscript, we isolate sperm from the cauda epididymis via tissue dissection and sperm swim out, and we then wash sperm with a somatic cell lysis buffer. After washing, purified cauda sperm exhibit no detectable (<0.1%) contamination by intact somatic cells (**Fig. S1A**, left panel). Although our original goal was to exactly reproduce the conditions used in Jung *et al* 2017 (Jung et al. 2017), we noticed that the investigators in this study did not add any reducing agent to properly lyse sperm. This was of concern as sperm are subject to extensive disulfide crosslinking during maturation, and failure to reverse these crosslinks via treatment with a reducing agent (DTT, β-ME, or TCEP) results in significant difficulties in recovering genomic DNA from sperm. This is illustrated in **Figs. S1B-C** – even after overnight SDS and proteinase K treatment, scant genomic DNA (∼5% of the levels recovered from DTT-treated sperm) is recovered in the absence of reducing agent. Moreover, microscopic examination of untreated sperm revealed abundant intact sperm heads even after extensive incubation in detergent and proteinase K, whereas DTT-treated samples were entirely lysed following detergent and proteinase K treatment (**Fig. S1A**, right panels). Taken together, these findings suggest the concerning possibility that many assays of sperm chromatin architecture – including but not limited to those using inadequate levels (**Fig. S1C**) of reducing agent to allow access to the sperm genome – could be contaminated by cell-free chromatin.

To test the hypothesis that published assays of sperm chromatin architecture might be contaminated by cell-free chromatin, we set out to characterize the impact of several treatments on sperm ATAC-Seq profiles. Four conditions were considered. First, purified and washed cauda sperm were subjected to the exact same ATAC-Seq protocol reported in Jung *et al* 2017 and used in several additional studies (Jung et al. 2017; Jung et al. 2019; Gou et al. 2020): we refer to this as the “untreated” case; given that no reducing agent was used to help permeabilize sperm, we predict that the resulting ATAC-Seq profiles should primarily – indeed, can *only* – reflect the organization of cell-free chromatin. Second, we pre-treat sperm with DNase I to eliminate any cell-free chromatin prior to probing with Tn5; we predict that little to no genomic DNA should be available for transposition after removing cell-free chromatin but without properly opening sperm. Third, we add appropriate levels (50 mM) of DTT prior to Tn5 treatment to allow access to the sperm genome; under these conditions we anticipate a mixed profile resulting from both cell-free chromatin and the bona fide sperm genome. Finally, by pretreating sperm with DNase I, then adding DTT prior to Tn5 probing, we hope to reveal the landscape of accessible chromatin in mature spermatozoa without any confounding contamination. Importantly, under these conditions we find no evidence that residual DNase activity subsequently affects sperm genomic DNA (**Fig. S1D** and below).

### Sperm ATAC-Seq profiles are sensitive to cell-free chromatin contamination

Our initial efforts to probe sperm chromatin architecture were carried out using the same “untreated” ATAC-Seq conditions reported by Jung *et al* (Jung et al. 2017). Under these conditions, we robustly reproduce the published ATAC-seq landscapes published in several prior reports (**Fig. 1A**, **Fig. 2, Figs. S2A-C**), as expected. Analysis of library insert lengths revealed the typical pattern observed in somatic cell ATAC-Seq libraries, with a ∼75 bp peak (after accounting for sequencing adaptor lengths) corresponding to paired Tn5 insertions into open regulatory elements, followed by a series of peaks every ∼150 bp corresponding to insertions flanking mono/di/trinucleosomes (**Fig. 1B, Fig. S2D**).

**Figure 1.**
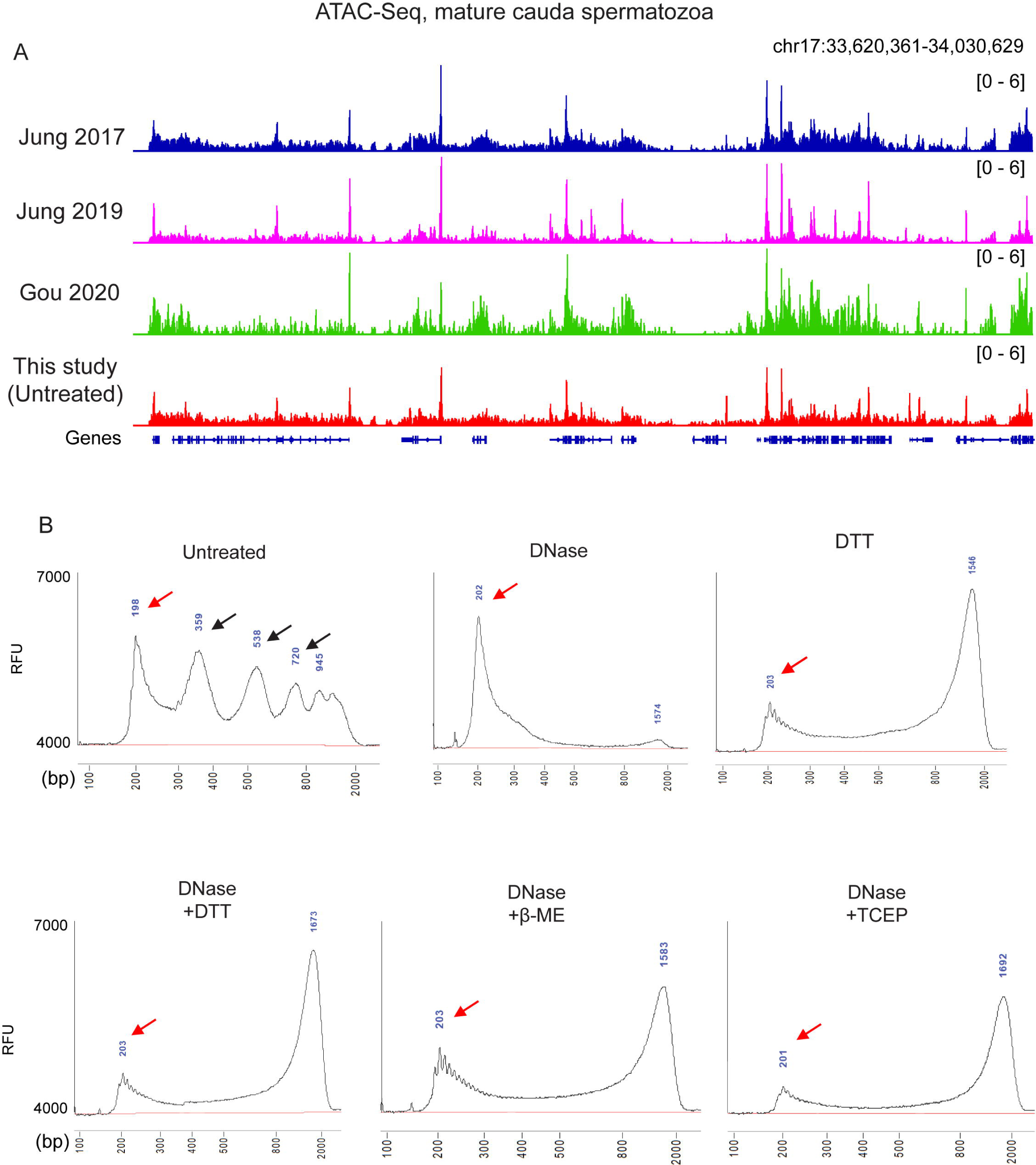
ATAC-Seq profiles from sperm subject following DNase and DTT treatment. A) Genome browser tracks for ATAC-Seq data from untreated sperm (this study) as well as three published ATAC-Seq datasets for sperm (Jung, et al. 2017; Jung et al. 2019; Gou et al. 2020). B) Bioanalyzer traces of ATAC-Seq sequencing libraries prepared from sperm samples pre-treated with the indicated conditions prior to Tn5 transposition. Red arrows show ∼75 bp inserts (after accounting for sequencing adaptors) corresponding to open chromatin regions, while black arrows in “untreated” libraries show insert lengths corresponding to mono-, di-, and tri-nucleosome length inserts. Notable in all the DTT-treated libraries is a peak at ∼1.3 kb. This peak reflects a landscape of random insertions across the sperm genome, with the size of the insert being an artifact of the PCR conditions used in library preparation. We confirmed this in two ways. First, we built a second set of ATAC-Seq libraries where the PCR extension time was extended to two minutes as compared to one minute in the libraries shown. Under these conditions, we find a ∼2.5 kb peak, confirming that PCR extension time is responsible for this peak location. Second, we gel-isolated the 1.3 kb material from a DNase + DTT ATAC-Seq library, sheared the DNA and characterized the material by deep sequencing. Resulting sequencing reads revealed a genome-wide background (not shown). (RFU: relative fluorescence units)

**Figure 2.**
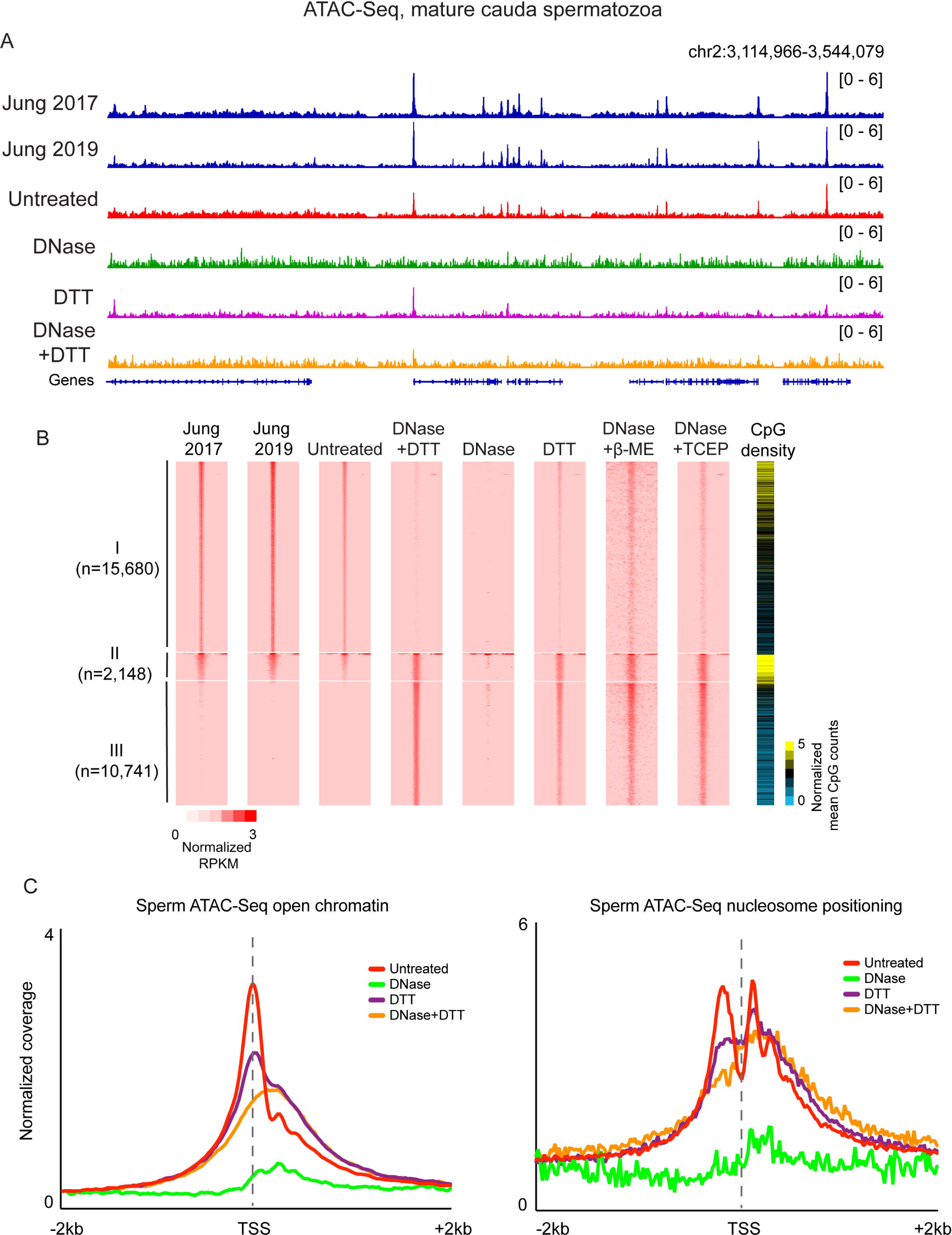
Published ATAC-Seq profiles in sperm are dominated by cell-free chromatin contamination. A) ATAC-Seq browser tracks for sperm subject to the indicated conditions. B) Heatmaps for the four indicated libraries, sorted into three categories: peaks specific to the untreated dataset (I; n= 13,605), shared peaks (II; n= 2,148), and peaks specific to DNase/DTT datasets (III; n= 8,627). C) Metagenes for <120 bp inserts – diagnostic of open chromatin – or for >150 bp inserts – diagnostic of nucleosome footprints – aligned over all annotated TSSs.

Turning next to sperm samples pre-treated with DNase I to remove cell-free chromatin prior to Tn5 probing, we generally failed to recover enough transposed DNA to build a sequencing library (not shown). When we did obtain sequencing libraries we recovered very low yields, with <10% of the levels of amplified DNA that could be recovered in the untreated condition (**Fig. S2E**). Analysis of insert lengths revealed a complete loss of the nucleosome signature (**Fig. 1B**), and sequencing profiles from these libraries were almost entirely flat; almost all the ATAC-seq peaks recovered in untreated sperm were lost after DNase treatment (**Fig. 2A-B**, **Fig. S2F-G**). These findings are consistent with the hypothesis that published ATAC-seq profiles generated from untreated sperm are most likely contaminated by cell-free chromatin.

### Eliminating cell-free chromatin reveals bona fide open chromatin profiles from mature sperm

We next turn to ATAC-Seq libraries prepared in the presence of the high levels of reducing agent required to enable access to Tn5 and other protein probes of chromatin architecture. We focus first on sperm preparations treated with both DNase and DTT. This pre-treatment should recover bona fide features of sperm chromatin, since it depletes cell-free chromatin before permeabilizing the sperm head. Analysis of library insert sizes revealed a dominant peak of ∼75 bp inserts (**Fig. 1B**), corresponding to short accessible loci, but no notable signature of nucleosome-sized footprints. This reflects a relatively flat landscape of Tn5 insertions across the primarily protamine-packaged genome, with any ∼150 bp nucleosomal footprints expected to result from low level of nucleosome retention (∼2%) in murine sperm being overwhelmed by this global background (albeit with potential implications for nucleosome positioning: see **Discussion**). Importantly, we find similar ATAC-Seq insert lengths in DNase-treated sperm that were permeabilized using other reducing agents such as β-ME or TCEP (**Fig. 1B**), further supporting our hypothesis that prior ATAC-Seq studies in murine sperm suffered from a failure to properly reverse disulfide crosslinks.

Comparing the DNase/DTT-treated ATAC-Seq landscape with the untreated landscape, we found that the majority of “untreated” ATAC peaks were lost and only a small fraction – associated with very high CpG-density regulatory elements – retained in DNase/DTT-treated sperm (**Figs. 2A-B**). Systematic peak calling identified 10,741 accessible loci in DNase/DTT-treated sperm samples, compared with 15,680 accessible loci in untreated samples, with 2,148 peaks shared between the two datasets at regulatory regions of unusually high CpG density. As in other cell types, ATAC peaks in sperm were overrepresented at promoters (**Fig. 2C**). However, the promoters that were uniquely identified in DNase-treated sperm were associated with reduced CpG density compared to the much higher CpG density found at the ATAC peaks from cell-free chromatin (**Fig. 2B**). Interestingly, we found no evidence for well-positioned 150 bp ATAC-Seq fragments – often considered diagnostic of nucleosome localization – surrounding accessible loci (**Fig. 2C**). This argues against a standard organization with accessible regulatory elements flanked by a pair of positioned nucleosomes; instead, we observe a modest but somewhat fuzzy/delocalized enrichment of 150 bp footprints across promoters (see **Discussion**). In terms of biological functions, accessible peaks in the DNase + DTT dataset were enriched near genes involved in transcriptional control and in early development, in contrast to various cell cycle and RNA splicing annotations specific to untreated sperm peaks (**Fig. S3**).

Finally, we consider ATAC-Seq libraries produced from DTT-only treated sperm. Consistent with these samples comprising a mixture of cell free chromatin and bona fide sperm chromatin, we found ATAC signal over both the peaks unique to untreated sperm as well as the DNase+DTT peaks (**Fig. 2**). This is true globally, with DTT-treated ATAC-Seq landscapes exhibiting a mix of both cell free accessibility peaks and bona fide sperm accessibility peaks (**Fig. 2B**). Moreover, metagene plots of open chromatin peaks and nucleosome signatures over transcription start sites revealed that DTT-only samples exhibited hybrid profiles between the untreated and DNase+DTT profiles (**Fig. 2C**), again supporting the idea that these libraries represent a mixture of cell free chromatin and bona fide sperm.

### DNase and DTT do not affect open chromatin in mESCs

Our findings in mature sperm support the hypothesis that prior ATAC-Seq studies of the sperm genome primarily reported on the status of cell-free chromatin, since this contamination is the only chromatin available in the absence of adequate reducing agent. However, we also considered the hypothesis that the DNase and DTT treatments might somehow drive artifactual changes to global chromatin architecture. To test this hypothesis, we performed ATAC-seq in a commonly-studied cell type – mESCs – treated with the same conditions used for sperm preparations. Our ATAC-Seq profiles from untreated mESCs broadly recapitulated known features of the ESC chromatin landscape (**Figs. 3A-B**), with widespread peaks of accessible chromatin, flanked by well-positioned nucleosomes, associated with promoters and enhancers (**Fig. 3C**). This landscape of open chromatin was almost entirely unaffected by DNase, DTT, or DNase + DTT treatments (**Fig. 3B and 3C**), confirming that these treatments do not grossly alter Tn5 transposition or chromatin organization in cell lines.

**Figure 3.**
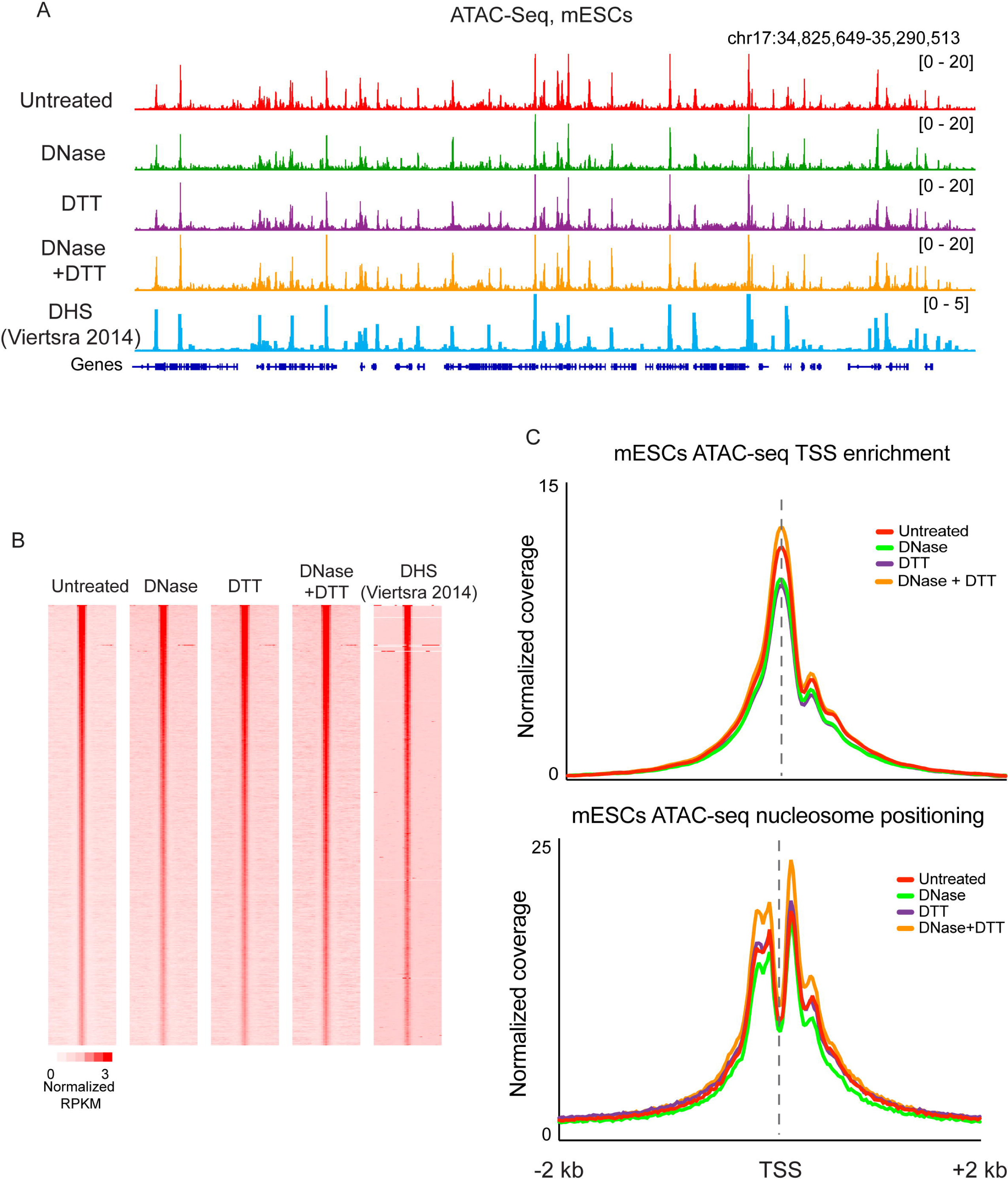
DNase and DTT treatment do not affect ATAC-Seq profiles in mESCs. A) ATAC-Seq profiles for mESCs treated with the indicated conditions, as well as published ESC DHS data from (Vierstra, et al. 2014). B) Heatmaps show ATAC read counts aligned over peaks (n= 58,130) called from untreated cells. C) Metagenes for <120 bp inserts, and >150 bp inserts, as in **Fig. 2C**.

### Contaminating chromatin likely derives from the epididymal epithelium

What is the origin of the cell-free chromatin that contaminates cauda epididymal sperm? Based on our previous efforts (Galan et al. 2021) focused on caput epididymal sperm – where the contaminating DNA skewed imprinting control regions (ICRs) from a germ cell profile (e.g., 0% or 100% methylated at ICRs) towards a somatic methylation profile (50% methylation) – we favor the hypothesis that contaminating DNA arises from somatic cells rather than cells of the germline lineage.

In order to test this hypothesis in cauda (rather than caput) epididymal sperm, we turned to NOME-Seq (Kelly et al. 2012), a protocol which leverages the bacterial M.CviPI methyltransferase to methylate accessible cytosines in GpC dinucleotides (rather than the endogenous CpG dinucleotide context). Importantly, the 5mC readout here enables simultaneous analysis of genomic accessibility at GpC dinucleotides, as well as endogenous cytosine methylation at CpG dinucleotides. We assayed cytosine methylation directly by Oxford Nanopore sequencing to avoid bisulfite-related fragmentation of the genome, which would prevent long-distance linkage between methylase accessibility and endogenous methylation on the same DNA molecule (Wang et al. 2019a; Shipony et al. 2020). This approach, known as Nano-NOME-Seq (Lee et al. 2020; Battaglia et al. 2022), provides simultaneous readouts of induced GC methylation and endogenous CG methylation on single DNA molecules with read lengths of tens of kilobases. This allows us to assess the developmental origin of cell-free DNA – soma vs germline – based on the endogenous CpG methylation program at imprinting control regions.

We subjected untreated (eg no DNase or DTT) cauda epididymal sperm to M.CviPI methylation, then isolated genomic DNA using DTT treatment to allow recovery of both bona fide sperm DNA along with any cell-free contamination, and assayed cytosine methylation genome-wide by Nanopore sequencing. This initial untargeted analysis covered the genome at ∼2-3X depth, revealing that roughly ∼5-10% of sequenced GpC dinucleotides were methylated, consistent with our prediction that only cell-free DNA would be available for M.CviPI methylation. However, despite the good length of the reads (median ∼18-25 kb) in this initial dataset, relatively few reads included the small number of ICRs where we could easily distinguish somatic from germline-derived genomes.

We therefore used CRISPR-Cas9 to target our Nanopore sequencing to more deeply sequence several ∼25-30 kb regions surrounding well-characterized ICRs (Xie et al. 2012; Gilpatrick et al. 2020). The resulting dataset was far deeper, with an average of 150 reads spanning each of the targeted ICRs (**Figs. 4A-B**). In order to assess the lineage of origin for cell-free DNA, we first separated reads according to the extent of methylation at GC dinucleotides to separately analyze methylase-accessible reads (with average GpC methylation >=20%; reporting on cell-free DNA with a total of 26,729 reads) and methylase-protected reads (with average GpC methylation < 20%; reporting on sperm with a total of 341,117 reads). The estimated cell-free DNA contamination was thus 7.8% here. As expected of methylation in the germline, ICRs in sperm – defined based on their protection from M.CviPI methylation – were either fully CpG-methylated (paternally imprinted) or fully CpG-unmethylated (maternally imprinted) (**Figs. 4A-B**). In contrast, accessible – eg, GC-methylated – genomic reads exhibited mixed CpG methylation profiles (∼50%) at ICRs (**Figs. 4A-B**), confirming our prediction that cell-free DNA is derived from somatic cells rather than sperm. Together, these data provide independent validation of our hypothesis that protein probes (Tn5, M.CviPI, antibodies, MNase, etc.) cannot access the genome of sperm that have not been permeabilized by DTT treatment, and support our prediction that this accessible DNA/chromatin is somatic in origin.

**Figure 4.**
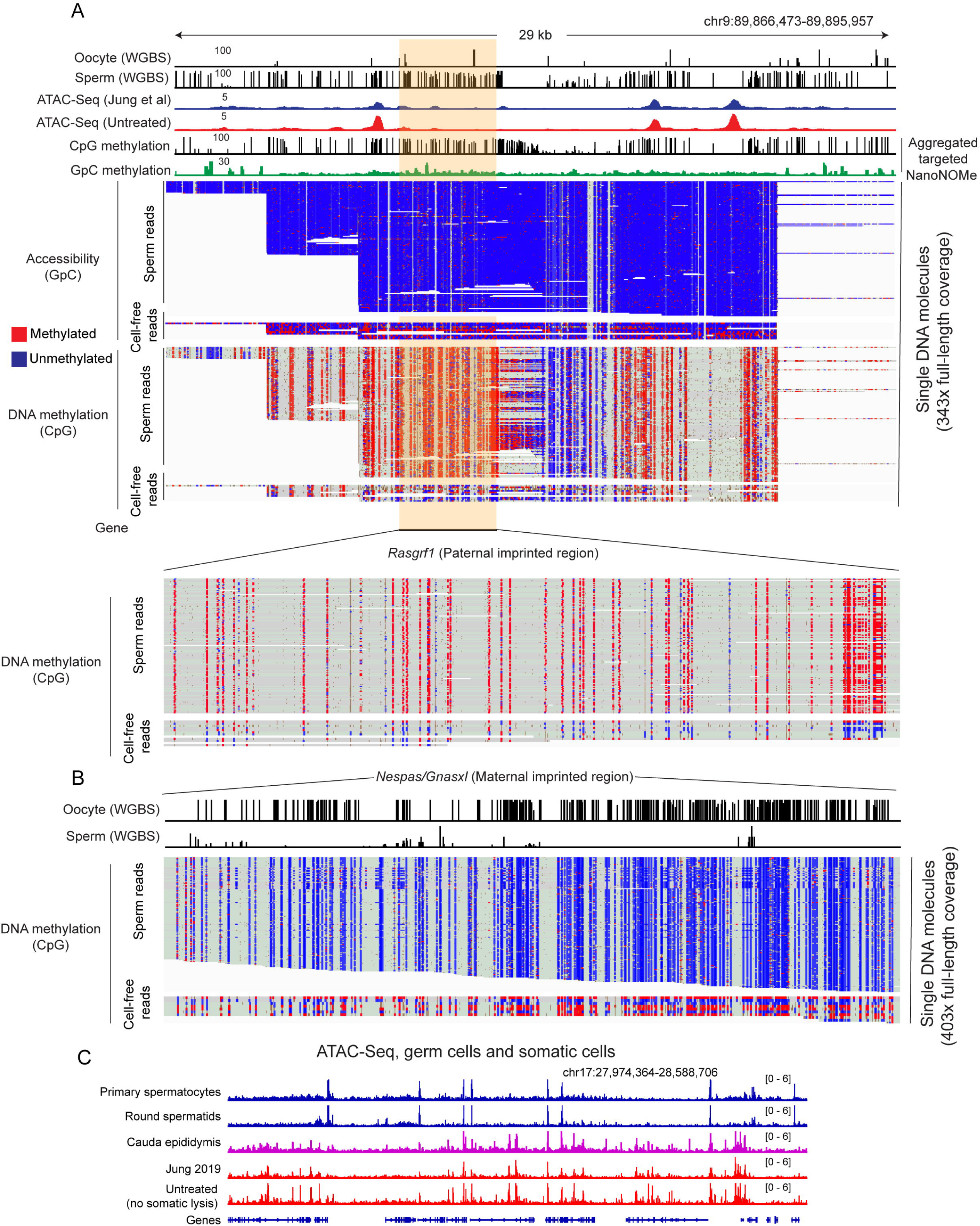
Contaminating cell-free DNA is of somatic origin. A-B) Nano-NOME-Seq data for two imprinting control regions in untreated sperm. Untreated sperm were subject to M.CviPI-driven GpC methylation, then genomic DNA was extracted after DTT treatment to obtain DNA from both contaminating material as well as sperm, and resulting DNA was subject to long-read sequencing by Oxford Nanopore. Resulting methylation calls (red and blue represent methylated and unmethylated cytosine, respectively) are shown separately for CpG methylation and GpC methylation (analysis was restricted to HCG and GCH – where H represents A/C/T – to avoid ambiguous GCG methylation), as indicated. In addition, reads are separated based on the extent of GpC methylation – the majority of reads exhibit low (<10% GpC methylation), whereas a small fraction of reads exhibit >20% GpC methylation and thereby represent accessible DNA molecules in untreated sperm preparations. Importantly, DNA molecules protected from M.CviPI – arising from the bona fide sperm genome – exhibit the 0% or 100% methylation expected at imprinting control regions in germline samples. In contrast, the small fraction of GC-methylated reads – reflecting accessible cell-free chromatin – include a mix of methylated and unmethylated ICRs consistent with a somatic origin for these DNA molecules. C) ATAC-Seq data for the indicated samples, including ATAC-seq for cauda epididymal epithelium. Key here is the strong agreement between ATAC profiles for untreated sperm (representing contaminating chromatin) and cauda epididymal epithelium.

Beyond this, our methylation-based study cannot more precisely define the cell of origin for contaminating DNA in cauda epididymal sperm preparations, in part because to our knowledge there have been no methylation surveys of relevant reproductive support cell types (e.g., Sertoli cells in the testis, principal cells in the epididymis, etc.). That said, comparing ATAC-Seq profiles from sperm isolated following relatively gentle vs. relatively disruptive dissection protocols (single slice followed by swim out vs. mincing and squeezing) revealed more extensive contamination resulting from more disruptive dissection (**Figs. S2A, D**). We therefore generated ATAC-Seq libraries from cauda epididymis epithelium. These profiles broadly agreed with both prior ATAC-Seq datasets (Jung et al. 2017; Jung et al. 2019; Gou et al. 2020), as well as our own dataset, from untreated sperm (**Fig. 4C, Fig. S2B**). Conversely, contaminated ATAC-Seq data – our untreated sperm data, or published datasets (Jung et al. 2017; Jung et al. 2019; Gou et al. 2020) – exhibited poor correlation with data generated from sperm precursors (spermatocytes and round spermatids (Maezawa et al. 2018): **Fig. 4C, Fig. S2B**). Taken together, these data definitively show that contamination of cauda epididymal sperm arises from somatic tissues, most likely epithelial cells of the epididymis.

### Contamination of histone modification profiles by cell-free chromatin

Our ATAC-Seq data provide strong evidence that epigenomic studies of untreated sperm report primarily on contaminating cell-free chromatin. This finding naturally raises questions about other published aspects of sperm chromatin architecture. Given that multiple studies have implicated sperm histone modifications in programming early gene expression in the preimplantation embryo (Hammoud et al. 2009; Brykczynska et al. 2010; Erkek et al. 2013; Jung et al. 2017; Jung et al. 2019; Lismer et al. 2020; Yoshida et al. 2020; Lismer et al. 2021), and perturbing histone modifications during spermatogenesis has phenotypic effects in the next generation (Siklenka et al. 2015; Lesch et al. 2019), we set out to explore whether the histone modification landscape of sperm might also be affected by cell-free chromatin in the epididymis.

We initially carried out CUT&RUN and CUT&Tag (Skene and Henikoff 2017; Kaya-Okur et al. 2019) to identify genomic loci associated with H3K4me3 or H3K27me3, histone modifications related to the Trithorax and Polycomb epigenetic memory systems (Schuettengruber et al. 2007) that have been mapped in sperm in multiple prior studies (Hammoud et al. 2009; Brykczynska et al. 2010; Erkek et al. 2013; Jung et al. 2017; Jung et al. 2019; Lismer et al. 2020; Yoshida et al. 2020; Lismer et al. 2021). In both cases, we found that libraries prepared from untreated sperm recapitulated modification landscapes previously obtained using ChIP-Seq (Erkek et al. 2013; Jung et al. 2017) (**Fig. S4**), but that these profiles were completely lost following DNase treatment. This again argues that prior histone modification maps were likely contaminated by cell-free chromatin. Curiously, we were unable to elicit antibody-specific CUT&RUN or CUT&Tag profiles from DTT-treated sperm (with or without DNase pretreatment – **Fig. S4**); we have thus far been unable to define conditions that yield antibody-specific profiles for these protocols.

We obtained similar results using ChIP-Seq (Johnson et al. 2007) on formaldehyde-fixed sperm chromatin sheared via sonication (**Fig. 5, Fig. S5, Methods**). We focused on a sequence-specific DNA-binding protein, CTCF, and the well-studied histone modifications H3K4me3 and H3K27me3 (K4/27), as these epitopes were expected to exhibit distinctive localization profiles. For all three epitopes, ChIP-Seq profiles from DTT-treated sperm samples – where we expected to see a mixture of contaminating chromatin and bona fide sperm – yielded profiles consistent with prior mapping studies (Erkek et al. 2013; Jung et al. 2017) (**Fig. 5A**). Peaks from the CTCF dataset were strongly enriched for the CTCF binding motif, whereas K4/27 profiles were associated with promoters and repressed developmental regulators, respectively (**Figs. 5B-C, Fig. S5A**).

**Figure 5.**
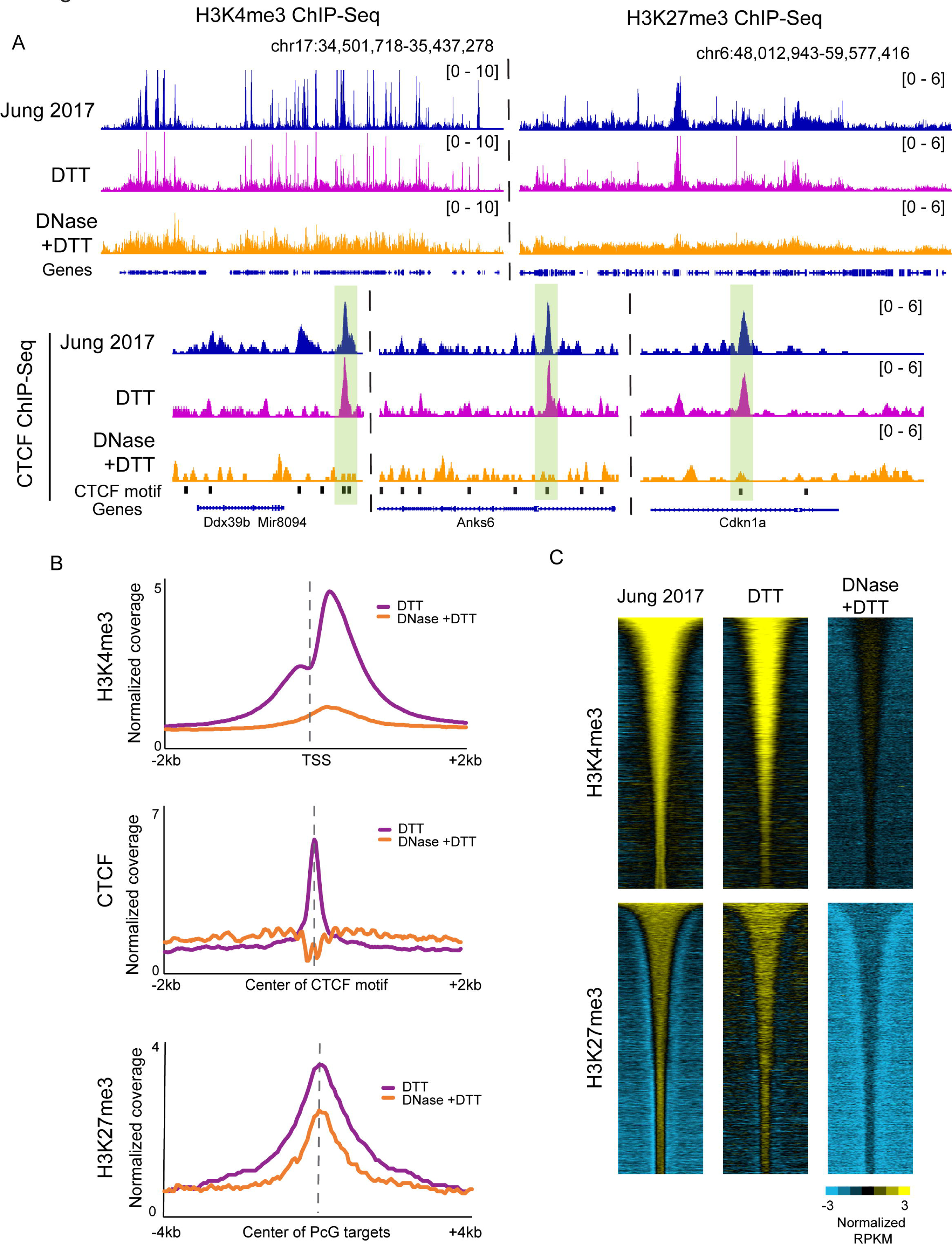
Histone modification profiling in DNase-treated sperm reveals a loss of specific signal. A) Browser tracks for H3K4me3, H3K27me3, and CTCF ChIP-Seq in sperm. In all three cases top panels show data from published sperm datasets (Jung, et al. 2017), along with our data from DTT- and DNase+DTT-treated sperm underneath. B) Metagenes aligned over the relevant peak locations (TSS, CTCF motif, and Polycomb-group (PcG) targets), showing H3K4me3, H3K27me3, and CTCF enrichment in our DTT only and DTT+DNase datasets. C) Heatmaps show H3K4me3 and K27me3 enrichment over all peaks from Jung *et al*. 2017, with data shown for Jung *et al* alongside our DTT and DTT+DNase datasets.

However, all three ChIP-Seq profiles were dramatically altered in maps generated from DNase+DTT-treated sperm (**Fig. 5A**). In the case of CTCF, we obtained a flat landscape, and those peaks that were called from this dataset were not enriched for the CTCF motif (**Fig. 5B, Fig. S5B**). This flat landscape suggests either that CTCF is not in fact associated with the mature sperm genome, or that our ChIP conditions – whether incomplete fragmentation and solubilization of the sperm genome (**Fig. S5C**), or fixation conditions that impact CTCF epitopes needed for immunoprecipitation – precluded accurate identification of CTCF localization. Similarly, DNase treatment also dramatically altered the localization landscape of K4/27. Significantly, the localization of these histone marks in DNase+DTT-treated sperm were distinct from the CTCF landscape; ChIP-Seq in bona fide sperm chromatin therefore exhibited somewhat antibody-specific profiles, unlike CUT&RUN or CUT&Tag (**Fig. S4**). Nonetheless, all three ChIP datasets revealed relatively flat landscapes – quite distinct from landscapes previously reported – with little evident biological significance.

As a fourth approach to exploring the histone modification landscape in sperm, we carried out native MNase ChIP-Seq (Liu et al. 2005; Erkek et al. 2013; Jung et al. 2017) for K4/27 methylation (**Fig. S5D**). Here, we again recover published K4/27 maps using untreated sperm; however, for this assay we observe no substantial changes following DNase pretreatment, with or without DTT-dependent permeabilization (**Fig. S5D**). At first glance, this finding might suggest that prior reports were in fact able to accurately measure sperm chromatin uncontaminated by cell-free chromatin contamination. However, given the dramatic effects of DNase treatment on the other three histone modification assays here, along with the inability of protein probes to access sperm in the absence of DTT treatment – raising the question of what is being mapped in the DNase-only MNase ChIP-Seq dataset – we instead speculate that some intact nucleosomes remain intact and available for ChIP following DNase treatment of the cell-free chromatin. Conversely, the destruction of adjacent linker DNA by DNase treatment would prevent Tn5 insertion adjacent to remaining nucleosomes, thus preventing specific ATAC or CUT&Tag readout (see **Discussion**).

Altogether, we successfully recover prior K4/27 landscapes from untreated sperm using four distinct assays: CUT&RUN, CUT&Tag, sonicated/fixed ChIP-Seq, and native MNase ChIP-Seq. However, each of these assays exhibits distinct behavior following DNase or DTT treatment; together, these findings raise substantial concerns regarding affinity-based chromatin analysis in bona fide sperm, and motivate continued optimization to allow more efficient recovery of proteins of interest.

### Contamination affects published chromosome folding maps from mature sperm

We finally turn to the question of how the sperm genome is folded in three-dimensional space, focusing on reports using the gold standard molecular approach to chromosome folding, Hi-C (Lafontaine et al. 2021). A number of studies have generated Hi-C maps for mature murine spermatozoa (Battulin et al. 2015; Jung et al. 2017; Ke et al. 2017; Jung et al. 2019; Vara et al. 2019; Wang et al. 2019b), with the majority of these studies (but see (Vara et al. 2019): discussed in the **Fig. S6** legend) reporting maps with typical features of somatic cells – A and B compartments corresponding to euchromatin and heterochromatin, and TADs with flares and loop anchors characteristic of loop extrusion processes. These maps are quite surprising given the compaction required to fit the genome into the small sperm nucleus, as well as the unusual physicochemical properties (Balhorn 1982; Balhorn 2007) of the major packaging proteins – the protamines – that organize the sperm genome. Notably, a review of methods sections of published sperm Hi-C papers revealed that the majority of these efforts neglected to add DTT or other reducing agents required to access the sperm genome (**Figs. S1B-C**), and are thus likely to assay only cell-free or other contaminating somatic cell chromatin architecture.

To test this hypothesis, we generated Hi-C maps for spermatozoa using DNase and/or DTT as in the various experiments detailed above. **Fig. 6** shows Hi-C contact maps for data from Jung *et al*. 2017 alongside our DTT only and DNase+DTT libraries. Data from Jung *et al* reveal the typical “checkerboard” pattern of A/B compartment organization at megabase scales (top panels), along with boxes on the diagonal with dots at the corners diagnostic of loop extrusion-mediated organization of gene-scale domains into TADs (middle panels). In our DNase+DTT samples all of these features are lost, with the exception of a faint signal of compartment organization (**Fig. 6**, bottom row). Importantly, the absence of TADs seen here has previously been reported in Hi-C studies of human sperm (Chen et al. 2019), as well as in Hi-C studies in both human and *Xenopus* sperm in the accompanying manuscript from Jessberger *et al*. We note that although this absence of specific organization could plausibly arise from certain types of chromosome packaging behavior, a more likely possibility for the limited structure seen in sperm Hi-C is that protamines are inefficiently crosslinked by formaldehyde and so standard Hi-C crosslinking conditions fail to capture the chromosomal contacts that do occur in the sperm nucleus (see **Discussion**).

**Figure 6.**
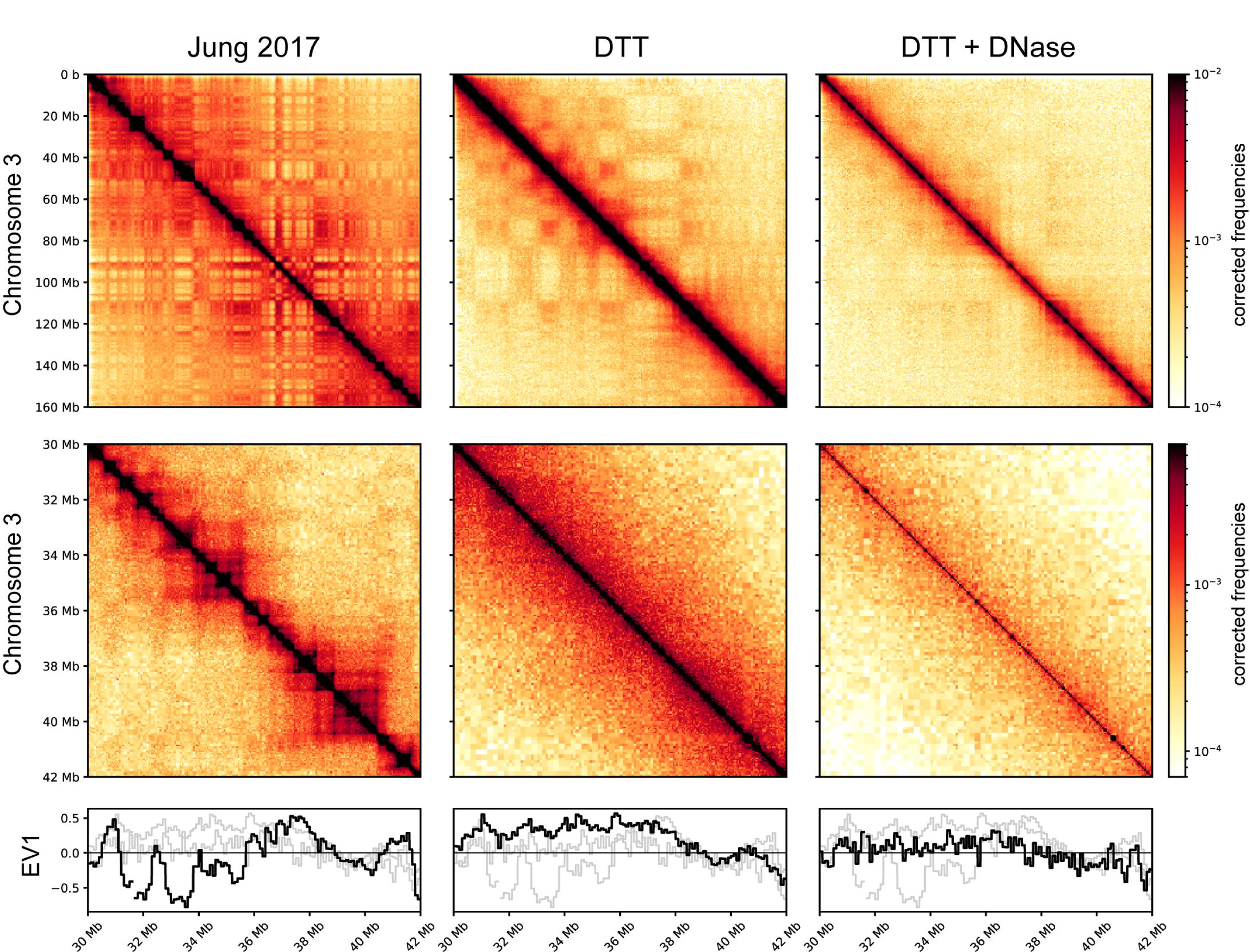
Absence of compartment signals and TADs in bona fide sperm Hi-C maps. Hi-C contact maps for (from left to right) untreated sperm (data from (Jung, et al. 2017)), DTT-treated sperm, and DNase and DTT-treated sperm. Top panels show a zoom-out genome view covering chromosome 3, middle panels show a typical ∼10 MB zoom in (Chr 3:30-42 MB). Bottom panels show A/B compartment calls (EV1: eigenvector 1) for the zoomed-in region.

Finally, as seen with our ATAC-Seq datasets, the DTT-only samples exhibit behavior consistent with a mixture between the “empty diagonal” of the DNase+DTT sample with a small contribution from the contaminating signal seen in untreated sperm libraries. This again fits our predictions and further supports our contention that even using DTT to break sperm nonetheless allows for contamination by cell-free chromatin if it has not been eliminated prior to assay.

## DISCUSSION

Here, we revisit the unusual genomic packaging state in mature mouse spermatozoa. Most importantly, we find that a wide variety of published analyses of sperm chromatin compaction are likely to have been contaminated by cell-free chromatin. We show that DNase I treatment of sperm prior to lysis removes this contaminating material, and further show that inadequate use of reducing agents in many studies precludes analysis of bona fide sperm chromatin by numerous commonly-used methods, including ATAC-seq, CUT&RUN, CUT&Tag, ChIP-Seq, and Hi-C. Together, our findings force a reappraisal of the packaging of the haploid genome in the compact sperm nucleus, and raise new questions about the potential role for sperm chromatin as a carrier of epigenetic information from fathers to offspring.

### Sperm chromatin assays are contaminated by cell-free chromatin

We previously identified cell-free chromatin as a major (∼20-30%) contaminant of immature sperm from the caput epididymis (Galan et al. 2021; Chen et al. 2022) based on cytosine methylation data. Definitive demonstration that unexpected cytosine methylation patterns resulted from cell free DNA was obtained by showing that pretreating caput sperm with DNase was sufficient to restore cytosine methylation to the levels seen in other germ cell populations. Although we did not highlight potential contamination of cauda sperm, even in this population we found that DNase treatment resulted in a ∼2-5% reduction in methylation at sites that are hypermethylated in cell-free chromatin compared to sperm (Galan et al. 2021). Further evidence for cell-free DNA contamination in cauda sperm can be seen in attempts to isolate sperm genomic DNA with and without DTT (**Figs. S1B-C**) – in the absence of DTT, relatively little DNA is obtained even after Proteinase K and SDS treatment overnight, with the yield of this DNA being ∼5% of the DNA obtained when sperm are properly broken using reducing agents.

The origin of this cell free chromatin is unclear – in the *caput* epididymis, it is associated with citrullinated histones (Galan et al. 2021) and thus may arise in situ from NETosis (Sollberger et al. 2018) or a similar programmed cell death process. It is worth noting that cell free chromatin in the caput epididymis co-purifies with sperm isolated by FACS (Chen et al. 2022); it is therefore clear that in the absence of enzymatic removal, somatic DNA is likely to contaminate murine sperm preparations no matter how extensive the purification. As seen for caput epididymal sperm, contamination of *cauda* epididymal sperm is also clearly derived from somatic cells (**Figs. 4A-B**), rather than a small population of easily-lysed or otherwise defective sperm populations. Importantly, contamination in the cauda epididymis varies depending on the extent of tissue disruption during dissection (**Methods**; **Figs. S2A, D**), and thus could partly or entirely arise during this process. Consistent with this, ATAC-Seq profiles from untreated sperm are best-correlated with ATAC data generated from the epididymal epithelium (**Fig. 4C, Fig. S2B**). Nonetheless, we still observed artifactual ATAC-Seq signal even in sperm obtained using the gentlest dissections (**Figs. S2A, D**), indicating that contaminating material – whether generated during the process of dissection, or already present in the epididymal lumen – is present no matter how gentle the dissection.

This subtle contamination is of little importance to studies of cytosine methylation (assuming sperm are obtained from a well-defined cauda epididymis dissection without substantial vas deferens), as extraordinarily deep sequencing is required to obtain precision greater than 2% in whole genome bisulfite sequencing. In contrast, studies of cauda sperm *chromatin* are particularly susceptible to this contamination owing to two idiosyncrasies of sperm biology. The first is the aforementioned requirement for a reducing agent for effective sperm lysis, driven by the extensive disulfide crosslinking that occurs during sperm maturation (Calvin and Bedford 1971; Saowaros and Panyim 1979). Thus, any studies of sperm chromatin carried out in the absence of DTT or other reducing agents – including the majority of sperm Hi-C studies as well as a substantial number of 1-dimensional chromatin studies – can only possibly survey the landscape of contaminating chromatin. The second challenge, affecting even studies performed using DTT, is that the vast majority of nucleosomes are evicted during spermatogenesis, leaving a small population (∼2% in mouse) of nucleosomes retained in mature spermatozoa. Thus, contamination by ∼2-5% cell free chromatin, presumably completely nucleosome-associated or nearly so (**Fig. S2F**), is sufficient to compete with or even overwhelm any true signal from sperm chromatin. This is of course even more problematic in scenarios where the bona fide sperm genome cannot be assayed for other technical reasons, as may occur for Hi-C (see below).

Our ATAC-Seq data confirm precisely our predictions based on the considerations above. In the absence of DNase or DTT, we robustly recapitulate prior ATAC-Seq studies performed under these same conditions (**Fig. 1**). Our contention is that these accessibility peaks reflect cell-free chromatin, a hypothesis that is further supported by the nearly complete loss of signal obtained when using sperm samples pre-treated with DNase I or MNase. Perhaps most compelling are the results from samples treated with DTT only, where we confirm the prediction that ATAC-Seq libraries should reveal a superposition of the cell-free chromatin peaks and the bona fide sperm peaks identified in DNase+DTT samples. Altogether our data are completely and parsimoniously explained by our hypothesis that sperm chromatin assays generated from untreated sperm samples can only capture aspects of cell free chromatin contamination.

### An updated view of genomic accessibility in murine sperm

Considering our ATAC-Seq data that was generated from properly broken sperm, free of cell-free chromatin, we found thousands of peaks of genomic accessibility that were only observed in DTT-treated sperm but not in cell-free chromatin (**Fig. 2B**). These peaks were enriched at regulatory elements associated with genes involved in transcriptional control, early development, and neurogenesis (**Fig. S3**). Whether the chromatin accessibility here is related to prior reports of H3K27me3 marking genes “poised” for early embryonic transcription (Hammoud et al. 2009; Brykczynska et al. 2010) bears further exploration; the difficulty of recovering specific histone modification profiles from uncontaminated sperm (**Fig. 5**, **Figs. S4-5**) precludes us from drawing any clear conclusions. In any case, this landscape of chromatin accessibility is of course completely distinct from the ATAC-Seq profiles of contaminating cell-free chromatin (**Figs. 1-2**) which are enriched for cell cycle and RNA splicing genes (**Fig. S3**).

Our data also provide some indirect evidence regarding nucleosome organization in mature sperm. Specifically, ATAC-Seq libraries not only capture relatively accessible genomic loci as relatively short ∼75 bp DNA fragments, but also provide information on nucleosomes flanking accessible regions (Buenrostro et al. 2013). These ∼140-150 bp fragments result from a pair of Tn5 transposition events, with one event occurring in a relatively open genomic locus along with a second insertion occurring at the far end of an adjacent nucleosome. Indeed, ∼140-150 bp ATAC peaks have been used in several studies of mammalian sperm to infer the existence of nucleosomes flanking regulatory elements (CpG islands, etc.) (Jung et al. 2017; Jung et al. 2019). However, in our DNase+DTT ATAC-Seq libraries we find no discernable ∼150 bp peak in the insert length distribution (**Fig. 1B**), and analysis of mapped 150 bp footprints reveals diffuse enrichment rather than any evidence for positioned peaks surrounding accessible genomic loci (**Fig. 2C**). These data are not evidence against the existence of nucleosomes in sperm – nucleosomes not adjacent to open chromatin require two Tn5 insertions into short linkers and are heavily underrepresented in ATAC-Seq libraries of somatic cells – but instead suggest that any sperm-retained nucleosomes may not be highly enriched surrounding accessible regulatory elements. That said, it is clear that the location of MNase footprints are heavily dependent on technical variables from fixation to MNase digestion extent, and it is likely that the same will hold true for ATAC-based accessibility metrics, raising the difficult question of how to independently substantiate the packaging of the sperm genome inferred from assays using large molecular probes.

Together, these considerations raise the question of the molecular nature of accessible genomic loci in mature sperm. Are these regions relatively depleted of protamines? Are they occupied by other proteins, or broadly protein-depleted? Given our concerns with affinity-based chromatin assays in sperm (below), answering these questions in molecular detail will be challenging and will require optimization of affinity-based chromatin assays suited to mature sperm.

### Absence of compartments and TADs in mature murine sperm

We next consider the three-dimensional folding of the sperm genome, where multiple previous studies have reported the a priori unlikely finding that the sperm genome is organized nearly-identically to the genomes of fibroblasts and ES cells. As noted in the **Introduction**, all of the studies making this claim were performed in the absence of sufficient reducing agent, and thus can *only* represent cell-free chromatin. Consistent with this model, we note that multiple studies (Alavattam et al. 2019; Patel et al. 2019; Wang et al. 2019b) have explored dynamic changes in chromosome folding during spermatogenesis, reporting that typical architectural features – compartments, TADs, loops – are present in spermatogonia and progressively lost during sperm development, only to be inexplicably regained in mature spermatozoa and then immediately lost in the paternal pronucleus of the zygote. These data are consistent with dramatic alterations to chromosome folding occurring during spermatogenesis, combined with a failure to appropriately measure the chromosome folding state in mature spermatozoa due to contaminating extracellular chromatin.

Indeed, as with all the other assays of sperm chromatin, we find that the organization of the sperm genome into compartments, TADs, and loops, cannot be reproduced when sperm are properly broken (**Fig. 6**). We instead find an essentially featureless diagonal in the DNase+DTT condition. Importantly, here again – as with ATAC-Seq – we show that DTT-only datasets exhibit a mixture of the bona fide sperm profile with the cell free chromatin profile, which provides compelling evidence for our overarching hypothesis.

The featureless diagonal seen in our DNase+DTT samples has two potential explanations. The first, which we do not favor, is that the sperm genome is compacted without any consistent architectural features, perhaps by protamine-mediated aggregation/liquid-liquid phase separation as observed in vitro (Moritz et al. 2023). Alternatively, we speculate that standard Hi-C methodology fails to effectively assay interactions between genomic loci in physical proximity in sperm. We suspect that this results from the requirement for formaldehyde crosslinking to capture interactions between genomic loci in most 3C-derived protocols. Formaldehyde reacts efficiently with primary amines including the ε-amine moiety of lysine as well as primary amine groups on the nucleotide bases. Formaldehyde is expected to react much more inefficiently with the electron pair on arginine thanks to higher pKa (∼12.5 vs 10) of the guanidino group making this residue much less nucleophilic. Thus, while formaldehyde very effectively crosslinks DNA bases to histone lysines, and histone lysines to one another, in the case of protamines – which are highly enriched for arginine and in many species are completely devoid of lysines – formaldehyde may not capture protamine-protamine interactions. Indeed, the relatively flat interaction decay curves for our Hi-C data are consistent with poor crosslinking (**Fig. S6**).

We therefore consider our data extremely strong evidence arguing that prior sperm Hi-C studies in mammals have been contaminated by cell-free chromatin. Beyond this, at present we feel our data do not provide any insight into the three-dimensional organization of the mammalian sperm genome. For example, we find no evidence for the 50-80kb solenoidal structure seen in in vitro studies of protamine-DNA complexes, which would be readily visible in Hi-C as a second diagonal 50-80 kb away from the primary diagonal. Whether such a structure occurs in vivo will be of considerable interest for future studies. Productive avenues for future chromosome folding studies in sperm can be envisioned using specific arginine-reactive or DNA-reactive crosslinkers (Jones et al. 2019; You et al. 2021), crosslinking-independent chromosome folding assays such as Hi-TrAC (Liu et al. 2022) or GAM (Beagrie et al. 2017), or super-resolution microscopy “walks” to probe the organization of megabase-scale domains (Boettiger et al. 2016).

### Implications for sperm chromatin architecture studies in human

We note one intriguing finding that may provide important insights into differences between studies of human and mouse sperm chromatin. Briefly, Chen *et al* generated Hi-C maps from human sperm, finding a largely featureless diagonal like the one seen in our DNase+DTT dataset (Chen et al. 2019). The authors performed a valuable and insightful mixing experiment in which they performed Hi-C in a mixture of ejaculated human sperm and cauda epididymal mouse sperm, recovering the compartments and TADs reported in mouse sperm along with the flat diagonal they reported for human. Moreover, in the accompanying manuscript from Peters and colleagues, Hi-C studies of human and *Xenopus* sperm both failed to detect TADs, again highlighting the question of why human and mouse sperm differ so dramatically in their apparent chromosome folding behavior.

We consider two potential explanations for the discrepancy between human and mouse sperm Hi-C. The first is that it has been reported that human sperm are more readily permeabilized in the absence of DTT (Hisano et al. 2013), suggesting that under the reported conditions the mouse sperm genome was inaccessible and so mouse contacts could only be generated from cell-free chromatin, while – in contrast – the human sample was perhaps somewhat permeabilized (albeit capturing few bona fide chromosomal contacts thanks to poor fixation of protamines with formaldehyde), thereby yielding maps similar to our DTT-only data in **Fig. 6**. Alternatively, we believe it may be the case that ejaculated sperm are less contaminated by cell-free chromatin, as murine seminal fluid carries high levels of the DNase2b (Smyth et al. 2022), which could play a role in clearing any cell-free chromatin carried forward from the epididymis. In addition, at least some contaminating chromatin is likely to be produced during the process of epididymal dissection in mouse (**Fig. 4C, Figs. S2A, D**), again suggesting that ejaculated sperm samples obtained without tissue disruption might exhibit less cell-free chromatin contamination. Regardless of the explanation for the discrepancy between human and mouse Hi-C experiments, the fact that TADs are only observed in mouse sperm but not in human, along with the elimination of TADs in mouse sperm by removal of cell-free chromatin, together demonstrate that the mouse sperm genome is unlikely to be organized in TADs as previously reported.

### The state of histone modification landscapes in mature sperm

We finally discuss the implications of our study for the histone modification landscape in sperm, where we consider our findings less definitive. On the one hand, here again our data suggest that the prevailing view of sperm histone modifications – that nucleosomes flanking CpG islands carry the “bivalent” pair of H3K4me3 and H3K27me3 histone modifications (Erkek et al. 2013; Zheng et al. 2016; Jung et al. 2017) – might also reflect contamination by cell-free chromatin. Using four common affinity-based methods for histone modification mapping – ChIP-Seq (fixed/sonicated, and native/MNase), CUT&RUN, and CUT&Tag – in untreated sperm we readily reproduce the findings of multiple prior histone modification surveys (**Figs. 5 and S4-5**). For three of these assays, as in the case of ATAC-Seq, these patterns are dramatically altered when sperm are pre-treated with DNase I to eliminate cell-free chromatin, again consistent with extant histone modification profiles in sperm reflecting cell-free chromatin contamination. The only condition where we see no effect of DNase I treatment is for native MNase-ChIP-Seq, where unfixed sperm are digested with MNase prior to ChIP (**Fig. S5D**). We speculate that this results from some nucleosomes in the cell-free DNA remaining intact following DNase treatment – the absence of adjacent linkers in such material would prevent Tn5 insertion in ATAC or CUT&Tag methods, but resulting nucleosomes would remain available for immunoprecipitation. Whatever the explanation, the extraordinary differences in behavior for the four mapping approaches used here – CUT&RUN, CUT&Tag, ChIP-Seq from fixed and sonicated chromatin, and ChIP-Seq from native MNase-digested chromatin – raise significant concerns regarding extant histone modification mapping efforts in sperm.

That said, three issues prevent us from confidently rejecting the prevailing model of “bivalent” nucleosomes present at developmental promoters. Firstly, K4/27 bivalent domains at CpG islands have been reported not only in mouse but also in human sperm studies (Hammoud et al. 2009). As noted above, Hi-C studies diverge dramatically between these two species; if the explanation for this difference is that ejaculated sperm are not contaminated by cell free DNA, this would suggest that K4/27-marked CpG islands might be a bona fide feature of uncontaminated mature spermatozoa in human, and the similar findings in mouse would thus support this view of mammalian sperm chromatin. Secondly, analysis of histone modifications during fetal germ cell development and spermatogenesis in the testis revealed that a subset (47 of 92 promoters) of K4/27-marked poised genes overlap with those reported in mature sperm, suggesting a progressive shaping of histone modifications patterns towards the landscape reported for mature sperm (Lesch et al. 2013). As DNase treatment is commonly performed during testicular dissociation into single cells (thus preventing cfDNA contamination), this again might support the prevailing CGI/bivalency model.

That said, none of the testicular populations analyzed in that study will have completed the histone to protamine transition, so these data could be compatible with either model – after all, K4/27 domains occur at CpG islands in other cell types and thus could occur both in spermatocytes/spermatids prior to the histone to protamine transition, as well as in the cells giving rise to contaminating cfDNA, but not in mature sperm. Finally, unlike the alternative set of peaks identified for ATAC-Seq in DNase-treated sperm, we cannot be confident of a putative bona fide sperm histone modification landscape as we have yet to identify conditions that allow for reliable affinity-based chromatin localization studies in sperm, having obtained relatively flat or nonspecific landscapes for ChIP-Seq, CUT&RUN, and CUT&Tag using several antibodies.

Taken together, our results should motivate some skepticism regarding the prevailing model for the sperm chromatin landscape, especially given the dramatic changes to modification localization following DNase treatment, but this model may yet prove correct. Further optimization will therefore be required to properly assess protein localization across the sperm genome using antibody-based approaches.

## ACKNOWLEDGEMENTS

We thank N. Krietenstein for invaluable technical advice on sperm lysis conditions and Hi-C, B. Bernstein and W. Flavahan for helpful advice on Nano-NOME-Seq experiment design, and S. Hammoud, B. Lesch, and A. Peters for discussions and insightful comments on sperm chromatin architecture. Work is supported by NIH R01AG073238 (OJR), R01HD072122 (TF), R01HG003143 (JD), and R35GM128782 (XZL). J.D. is an investigator of the Howard Hughes Medical Institute.

## MATERIALS AND METHODS

### Ethics statement

Animal husbandry and experimentation was reviewed, approved, and monitored under the University of Massachusetts Medical School Institutional Animal Care and Use Committee (Protocol ID: A-1833-18).

### Mouse husbandry and tissue collection

All samples were obtained from male mice of the C57BL/6J strain background, consuming control diet Ain-93g, euthanized at 19 weeks of age.

### Cauda sperm isolation and purification

For most assays, we used Prep 1 (see **Fig. S2**): Cauda epididymis was briefly dissected in PBS to remove fat tissues. Cauda tissues were transferred to Donners medium and sperm were released by several cuts and gentle squeeze of the cauda tissues. The cauda was discarded, and medium was transferred to a 1.5 ml tube. Sperm swim-up was conducted at 37°C for 1hr. Next, the upper ∼1.3 ml medium was recovered and filtered through a 40 μm strainer. Sperm were pelleted, washed with PBS, and resuspend/incubated in somatic lysis buffer (0.01% SDS, 0.005% Triton X-100) for 10 min on ice. Purified sperm were washed with PBS and subjected to downstream treatment; For Prep 2, based on (Chen et al. 2021), cauda epididymis was punctured by a needle and the sperm were squeezed out using two forceps. The sperm clot was collected in 250 ul of Donners medium in a 1.5 mL tube, followed by gently adding 1 mL of Donners medium on top. Sperm swim-up was conducted at 37°C for 1hr and the upper ∼1 ml medium was recovered for experiments; For Prep 3, based on (Hisano et al. 2013), cauda epididymis was cut 4-5 times and directly put in 1 ml of Donners medium in a round-bottom falcon tube. Then, 4 mL Donner medium was added along the wall of tube and sperm were allowed to swim up at 37°C for 1hr. Next, the upper ∼4 ml of medium was recovered for sperm experiments.

### Cauda epididymis cell suspension

Cauda epididymis tissue was placed in PBS and cut into small pieces. Sperm were squeezed out as much as possible followed by 10 min incubation at 37 °C to further allow sperm swim-out. Only big chunk of epithelium tissues were recovered through a 200um cell strainer and washed with in IMDM+DNase media (Thermo Fisher 31980030; Sigma 11097113001). The chopped tissues were further digested with complete media (10% FBS in IMDM), ECM (Extracellular Matrix Digestion media), and IMDM+DNase for 20 min followed by 0.25% Trypsin, 50 ug/mL DNase for another 15 min. Reaction was neutralized by FBS and single cell suspension was washed with PBS and quality-checked under a microscope.

### Nuclease and DTT treatment of cauda sperm

Purified sperm pellet was resuspended in PBS. For DNase treatment, RDD buffer and DNase I (Qiagen 79254) was added at 1:10 and 1:40 (∼0.1 unit per μl) dilution in 1x PBS, respectively. DNase treatment was carried out at room temperature for 10 min; For MNase treatment, sperm were incubated in 1x PBS containing 85mM Tris-HCl pH 7.5, 3mM MgCl_2_, 2 mM CaCl_2_, 0.1 U/μl of MNase (Worthington, LS004798) for 5 min at 37 °C; For Exo_V treatment, sperm were incubated in 1x PBS containing 1x NEBuffer^TM^ 4, 1mM ATP, 0.1 U/μl of RecBCD (NEB M0345) for 20 min at 37 °C. After washing with PBS, sperm were incubated in 1x PBS containing sperm were treated with 50 mM DTT (50 mM TCEP or 1% β-ME) in PBS for 1 hr at RT and quenched by 100 mM NEM (N-Ethylmaleimide). Pretreated sperm were pelleted and washed with PBS and subjected to ATAC-seq.

### ATAC-Seq

ATAC-seq was carried out using the Omni-ATAC-seq protocol (Corces et al. 2017) with several modifications. About 100,000 sperm were on-ice lysed with hypotonic lysis buffer (10 mM Tris-HCl pH 7.4, 10 mM NaCl, 3 mM MgCl2, 0.1% NP40, 0.1% Tween-20, and 0.01% digitonin) for 10 min. Sperm were then resuspended in Tagmentation buffer (10 mM TAPS-NaOH pH 8.5, 5 mM MgCl2, 10% DMF, 0.05% Digitonin) and tagmentation was conducted by adding Tn5 transposase (Diagenode, C01070012) at 1:20 and incubated at 37C for 30min. Reaction was stopped by SDS and proteinase K and incubated at 55°C overnight. SDS was quenched by tween-20 and library amplification was done with 1x NEBNext High-Fidelity PCR Master Mix containing 0.5 μM indexed primers using the following PCR conditions: 72°C for 5 min; 98°C for 3 min; and 14 cycles at 98°C for 15 s, 63°C for 30 s, and 72°C for 1 min. mESCs and cauda epididymis cells were used for ATAC-Seq and processed in the same way as sperm. Libraries were pair-end sequenced on an Illumina NextSeq500 platform.

### ATAC-Seq data analysis

Paired-end ATAC-seq reads were adapter-trimmed and aligned to the mm10 genome using Bowtie2 (Version 2.3.2) with the parameters: -t -q -N 1 -L 25 -X 2000 no-mixed no-discordant. All unmapped reads, non-uniquely mapped reads and PCR duplicates were removed. MACS2 (Zhang et al. 2008) was used for peak calling with parameters: -- nolambda --nomodel. Reads with insert size larger than 150 bp were separated and used for nucleosome positioning analysis. TSS enrichment analysis was done using Homer annotatePeaks (Heinz et al. 2010).

### Nano-NOME-Seq

Nano-NOME libraries were prepared as previously described (Battaglia et al. 2022). Briefly, about 2 million sperm were on-ice lysed with hypotonic lysis buffer (10 mM Tris-HCl pH 7.4, 10 mM NaCl, 3 mM MgCl2, 0.1% NP40, 0.1% Tween-20, and 0.01% digitonin) for 10 min. Sperm were pelleted down and resuspended in 500 ul of 1x GpC buffer containing 150 μl of 1 M sucrose, 1.5 μl of 32 mM S-adenosylmethionine (SAM; NEB, B9003) and 50 μl of M.CviPI (NEB, no. M0227L). The suspension was carefully mixed and incubated for 7.5 min at 37 °C, followed by a boost with an additional 25 μl of M.CviPI and 1.5 μl of SAM for 7.5 min. The reaction was stopped by the addition of 25 μl of 10% SDS, 5 μl of 0.5 M EDTA, 6 μl of 20mg/mL proteinase K, and 28 μl of 1M DTT followed by overnight incubation at 55 °C. The total sperm DNA was recovered by PCI and ethanol precipitation. For whole-genome nanopore sequencing, 1 ug of sperm DNA was used for library preparation using Oxford Nanopore Ligation-based library prep kit (ONT, SQK-LSK110). For target-enriched sequencing, single-stranded CRISPR– Cas9 DNA oligonucleotides (20 nt) were designed using the ChopChop v.3 designer tool (chopchop.rc.fas.har-vard.edu) and purchased from IDT. sgRNA were prepared using EnGen^@^sgRNA synthesis kit (NEB, no. E3322S). RNP complexes were assembled individually by combining 8 μl of nuclease-free water, 1.5 μl of NEBuffer r3.1 (NEB), 3 μl of 300 nM sgRNA, 1 μl of Cas9 Nuclease (NEB, no. M0386), followed by a 20-min incubation at 25 °C. The different RNP complexes were then pooled in a single tube. Per experiment, 3-4 μg of sperm gDNA was dephosphorylated with 10 μl of rSAP (NEB, no. M0371S) and 16 μl of 10× rCutSmart buffer (NEB, no. B6004S) for 30 min at 37 °C, then heat inactivated at 65 °C for 5 min. DNA was then purified through PCI and ethanol precipitation. The dephosphorylated DNA sample was then combined with the pool of RNP complexes in a 1:9 ratio. The reaction was incubated for 30 min at 37 °C to enable Cas9 cleavage. After digestion, Cas9 was inactivated by the addition of a 1/25 volume of 20 mg/mL proteinase K followed by a 15-min incubation at 55 °C. Cleaved and purified DNA was purified by PCI and ethanol precipitation and was subjected to dA-Tailing using Klenow Fragment (NEB, no. E6053). Reactions were incubated for 30 min at 37 °C and immediately subjected to adapter ligation as per the manual of Oxford Nanopore Ligation-based library prep kit (ONT, SQK-LSK110). All nanopore libraries were sequenced using MinION flow cell (ONT, R10.4.1).

### Nanopore Data Processing

Base calling was performed on the raw FAST5 files with Guppy (v.3.0.3 or v.5.0.11, ONT), using a configuration file for high-accuracy DNA base calling on an R10.4.1 pore at 450 bases s^−1^. The resulting reads were then mapped to the mm10 mouse reference genome without alternate contigs using minimap2 v.2.11 with default settings for alignment of nanopore reads (-x map-ont). Reads that mapped with a quality score <50 were then filtered out using samtools v.1.7. CpG and GpC methylation were simultaneously called on the remaining reads using nanopolish v.0.11.1. Average CpG methylation was calculated for each group of CpGs that did not contain any GCGs. To visualize the methylation patterns of the reads with IGV v.2.12.2, we modified the individual reads in the alignment (BAM) files. All cytosines called as unmethylated were converted to thymine to simulate bisulfite conversion. This was achieved using code adapted from the Timp Laboratory’s nanopore-methylation-utilities (https://github.com/ timplab/nanopore-methylation-utilities). Once these converted files were loaded in IGV, we were then able to visualize methylation using IGV’s bisulfite mode. Tiled data file tracks showing aggregated CpG methylation levels were generated with igvtools v.2.4.16.

### CUT&RUN

CUT&RUN libraries were prepared following the same protocol as previously described (Chen et al. 2019) with minor modifications. Specifically, 0.1% NP40, 0.1% Tween-20, and 0.05% digitonin was used in the antibody incubation buffer and wash buffer, and SPRI beads were used for DNA extraction after proteinase K treatment. For H3K27me3 CUT&RUN, a rabbit anti-H3K27me3 antibody (1:100; Diagenode, C15410195) was used. For H3K4me3 CUT&RUN, a rabbit anti-H3K4me3 antibody (1:100; Diagenode, C15410003) was used. All CUT&RUN libraries were sequenced on an Illumina NextSeq500 platform.

### CUT&Tag

CUT&Tag libraries were prepared following the same protocol as previously described (Kaya-Okur et al. 2019). Antibodies used were against H3K27me3 (1:50; ThermoFisher, MA5-11198), H3K4me3 (1:50; abcam, ab8580), Histone H3 (1:50; abcam, ab1791) and Serine 2 Phospho-Rpb1 CTD (1:50; Cell Signal, 13499S). All CUT&Tag libraries were sequenced on an Illumina NextSeq500 platform.

### ChIP-Seq

ChIP-Seq experiments were conducted following the protocol described in (Hisano et al. 2013; Chen et al. 2021) with some modifications. For fixed-sonication-based method, sperm were first fixed with 1% formaldehyde in PBS for 10 min at room temperature, followed by quenching with Glycine. Next, sperm were pre-treated with 50 mM DTT and 1 mg/ml heparin at 37°C for 2 hr. Sperm were then quenched with NEM and washed with PBS twice. After lysing sperm in complete buffer containing detergent, equal volume of 2x RIPA buffer was supplied to make a 1x RIPA condition. Chromatin shearing was performed using Covaris S220 platform with parameters: peak power 140 W, duty factor 5% and 200 Cycles/burst for a total of 30 min. For native-MNase-based method, sperm were directly lysed in complete buffer containing detergent and digested with 15 Unit of MNase (Worthington, LS004798) for 5 min at 37 °C. Reaction was stopped by EDTA and supplied with equal volume of 2x RIPA buffer. Sonicated or MNase-digested sperm chromatin were recovered from the supernatant by spinning at 10,000 x g for 10 min at 4°C, and were directly subjected to immunoprecipitation with corresponding antibodies (H3K4me3: Diagenode, C15410003; H3K27me3: Diagenode, C15410195; CTCF: Millipore 07-792). Library preparation was carried out using NEBNext Ultra II DNA Library Prep kit (NEB E7645). All ChIP-Seq libraries were sequenced on an Illumina NextSeq500 platform.

### Histone modification data analysis

For both CUT&RUN and ChIP-seq datasets, paired-end reads were trimmed using Trim Galore (version 0.4.5), followed by reads alignment to the mm10 genome using Bowtie2 (Version 2.3.2) with the parameters: -t -q -N 1 -L 25 -X 700 no-mixed no-discordant. PCR duplicates were removed using “MarkDuplicates” from Picard Tools (version 2.8.0) (https://broadinstitute.github.io/picard/). Only non-PCR duplicates and uniquely aligned reads (alignment records without “XS” tag) were used for downstream analyses. The RPKM values for each 100-bp bin were calculated following the formula “read counts/((bin_length/1000) × (total_reads/10^6^)). Peak calling for H3K4me3 and H3K27me3 datasets were done by MACS2 with parameters: --nolambda --nomodel, and --broad-cutoff 0.1 ---nolambda --nomodel, respectively.

### Hi-C library construction

The sperm library collection was made as described before (Lafontaine et al. 2021) with some modifications. Briefly, cells were fixed in 1% formaldehyde and after washing in PBS, left for 1 hour in 50 mM DTT. After quenching with 100mM NEM (N-ethylmaleimide) and washing in PBS, cells were incubated with DpnII overnight at 37°C. After restriction digestion, cells were incubated for 4 hours at 23°C with the Klenow DNA polymerase I to fill in the 5’ overhang and incorporate biotin-14-dATP followed by chromatin ligation for 4 hours at 16°C with T4 DNA ligase (Invitrogen). Then cells were left overnight in 400 ng/ml proteinase K and 0.5% SDS at 60°C overnight to reverse cross-linking. After DNA purification and precipitation, DNA was sonicated to 200-300 bp fragments and an additional size selection was done with Ampure beads. To repair DNA ends after sonication, T4 DNA polymerase (NEB) and Klenow DNA polymerase (NEB) were used together with T4 polynucleotide kinase (NEB). Biotinylated ligation products were pulled down using streptavidin beads (Invitrogen) followed by A-tailing and adaptor ligation for library construction. All Hi-C libraries were sequenced on an Illumina NextSeq500 platform.

### Hi-C data analysis

Hi-C data was analyzed using the Open Chromosome Collective suite (https://github.com/open2c). Briefly, 50bp paired-end fastq files were mapped to mm10 mouse reference genome using the *distiller-nf* (https://github.com/mirnylab/distiller-nf), invoking bwa mem to map fastq pairs in a single-side regime (-SP). Aligned reads were allowed a 1bp flexibility for removal of optical and PCR duplicates using *pairtools* (https://github.com/mirnylab/pairtools). After removal of identical positions and strand orientations, remaining valid pairs were binned into contact matrices of various resolutions using *cooler* (Abdennur and Mirny 2020). An iterative balancing procedure (Imakaev et al. 2012) was applied to all matrices, ignoring the first 2 diagonals to avoid short-range ligation artifacts at a given resolution, and bins with low coverage were removed using MADmax filter with default parameters. Resultant “.cool” contact matrices were used in downstream analyses using *cooltools* (https://github.com/mirnylab/cooltools) and uploaded to a HiGlass server (Kerpedjiev et al. 2018) via the Reservoir Genome Browser (https://resgen.io/).

### Contact probability (*P*(*s*)) plots & derivatives

We produced *P(s)* plots per chromosome arm as outlined in the “contacts vs distance” section of cooltools (https://cooltools.readthedocs.io/en/latest/notebooks/contacts_vs_distance.html). Briefly, mm10 chromosome sizes and arm locations were downloaded as ViewFrames from the UCSC database using Bioframe (https://bioframe.readthedocs.io/en/latest/index.html). For each arm, interaction counts (diagsum) were generated from a 1kb cooler file for each distance from the self-self diagonal. Counts for all chromosome arms were aggregated, logbin-smoothed (σ = 0.05) and *P*(*s*) curves were normalized for the total number of valid interactions in each data set. Corresponding derivative plots were generated using numpy gradient to estimate the slope from each *P*(*s*) plot.

### Compartment analyses

When present, compartmentalization is the strongest signal and will be represented by the first eigenvector (EV1). We performed eigenvector decomposition on observed-over-expected contact maps at 100kb resolution separated for each chromosomal arm using the cooltools package derived scripts (https://cooltools.readthedocs.io/en/latest/notebooks/compartments_and_saddles.html).

## SUPPLEMENTAL MATERIALS

**Supplemental Figures S1-S6**

### SUPPLEMENTAL FIGURE LEGENDS

**Supplemental Figure S1. DTT is required to lyse mature spermatozoa**

A) DIC Images of typical cauda epididymal sperm samples before and after processing for ATAC-Seq. Top panels show sperm processed without the use of DTT, revealing unbroken sperm heads even after overnight incubation in SDS and proteinase K. Bottom panels show efficient sperm lysis for DTT-treated sperm under the same conditions.

B) DTT is required to recover genomic DNA from sperm. Native PAGE gel (as used for all subsequent DNA electrophoresis) shows the gDNA of sperm isolated from different methods. Here and in **Fig. S2**, we compared sperm obtained from the cauda epididymis using three different preparation methods. In each case the cauda epididymis was removed via two gentle incisions at the junctions with the corpus and the vas deferens. Preparations differed by how the epididymis was further treated (**Methods**) – briefly, in Prep 1, epididymis was subject to multiple incisions and sperm were squeezed out, in Prep 2 the epididymis was punctured by needle and sperm were squeezed out, and in Prep 3 the epididymis was incised and sperm were allowed to swim out without any squeezing. In all three cases tissue was incubated in Donners medium at 37 C for 1 hr, after which the sperm-containing supernatant was carefully recovered. Sperm were then washed and either left untreated or incubated with 50 mM DTT for 1 hour, then quenched with NEM. Samples were then treated identically with lysis buffer and were incubated in SDS and proteinase K at 55°C for 16 hours, and lysed in Trizol for genomic DNA extraction.

C) Genomic DNA recovery following permeabilization with varying levels of DTT. Maximal DNA recovery is observed at 50 and 80 mM DTT, with somewhat lower gDNA recovery from the 20 mM DTT condition. Consistent with our estimates from cytosine methylation studies (Galan et al. 2021), contaminating cell-free DNA represents ∼5-6% of the DNA recovered from a fully (50 or 80 mM) DTT-permeabilized sperm prep.

D) Genomic DNA recovered from DTT-permeabilized sperm, with or without pre-treatment with DNase I. Notably, DNase I pretreatment does not impact the integrity of gDNA recovered from sperm.

**Supplemental Figure S2. Effects of sperm purification methods on chromatin accessibility landscapes**

A) Heatmaps showing ATAC-Seq enrichment for the three sperm preparations (see **Fig. S1B**), aligned over peaks from Jung *et al*. (along with Jung *et al* data shown in the leftmost panel). Importantly, enrichment for open chromatin at peaks reported by Jung *et al*, and a nucleosomal insert landscape, were observed for all three methods, but these features were stronger for the more disruptive tissue dissections. This suggests that at least some of the cell free chromatin contamination explored here may arise from the process of tissue disruption, but that this is unavoidable even under the gentlest dissection methods.

B) Correlation matrix between the indicated genome-wide datasets. RPKM was calculated for 2kb genome-wide bins and spearman correlations were analyzed in a paired-wise manner. Two large blocks of well-correlated datasets are apparent. The first shows strong correlation between ES cell ATAC-Seq datasets obtained either from untreated cells or from cells following DNase, DTT, or DNase+DTT treatments (see also **Fig. 3**). The other group of well-correlated datasets include public data from Jung et al (Jung et al. 2017; Jung et al. 2019), data from untreated sperm (this study), and, importantly, ATAC-Seq data from the cauda epididymal epithelium (see **Fig. 4**).

Moreover, we find that skipping the somatic cell lysis steps – as conducted in Jung *et al* – resulted in still better agreement between our “untreated” ATAC-Seq profile and published ATAC-Seq datasets. This suggests that contaminating chromatin is not exclusively produced as a result of somatic cell lysis, and that detergent washing helps to remove contaminating material, albeit inefficiently.

Data for immature sperm populations are distinct from ESCs and untreated sperm, while DTT-treated sperm (whether DNase-pretreated or not) cluster separately from immature sperm, untreated sperm, or ESCs. (Exo_V: Exonuclease V; DHS: DNase-Seq)

C) Scatterplots showing local ATAC-Seq enrichment at 1 kb surrounding all TSSs, for the indicated pairs of libraries. All three datasets from untreated sperm (this study and Jung 2017 and 2019) are highly-correlated, while data from DNase+DTT-treated sperm are distinct from any untreated samples. DTT-only sperm exhibits intermediate correlations with both untreated and DNase+DTT treated sperm samples.

D) Insert length distributions for the sperm preparations 2 and 3 (corresponding insert lengths for Prep 1 are shown in **Fig. 1B**).

E) ATAC-Seq library yields for sperm treated with the indicated conditions prior to Tn5 treatment. All yields (mean of DNA library yields from two replicates) from other conditions were normalized to untreated group (Prep 1).

F) Untreated sperm were treated with the indicated nucleases, then sperm were pelleted by centrifugation and genomic DNA was recovered from the supernatant and visualized by gel electrophoresis (left panel). Note the nucleosomal bands in the MNase-digested material, consistent with cell-free contamination by chromatin rather than naked DNA. Right panel visualizes sequencing libraries prepared from supernatant following the indicated treatments.

G) Deep sequencing of the sperm preps from panel (F). Bottom three panels show the supernatant material from (F), revealing that contaminating material arises from the entire genome, rather than specific loci. ATAC panels for the nuclease-treated samples were prepared using sperm pelleted following the indicated nuclease treatments, but not permeabilized with DTT. Note that no enrichment is seen for sperm treated with the endonucleases DNase I or MNase, while exonuclease treatment leaves cell-free DNA available for ATAC-Seq. Importantly, the continued presence of ATAC peaks in this material confirms that cell-free chromatin cosediments with sperm through a gentle centrifugation step.

**Supplemental Figure S3. Functional enrichment in sperm open chromatin**

A) Bar graphs show p values (expressed as −log10 of the p value) for various Gene Ontology categories enriched in genes found near the indicated sperm ATAC-Seq peaks.

B) Heatmap shows expression (in TPM) of genes located near DNase+DTT ATAC-Seq peaks at the indicated stages of spermatogenesis (Hammoud et al. 2014).

**Supplemental Figure S4. Lack of antibody specificity in sperm CUT&RUN and CUT&Tag profiles**

A) CUT&RUN profiles for the indicated histone modifications, for sperm either untreated or treated with DNase I and DTT. Published ChIP-Seq profiles from the indicated studies (Erkek et al. 2013; Jung et al. 2017) are shown above the CUT&RUN libraries for comparison.

B) Heatmaps showing H3K4me3 signal for the indicated libraries, aligned over peaks called from Erkek *et al* (Erkek et al. 2013)

C) Metagene showing averaged H3K4me3 enrichment for a 4 kb window surrounding all transcription start sites.

D) As in (A), for CUT&Tag libraries. Note that four distinct epitopes, including one which should be absent from mature sperm (Pol2S2P), all exhibit highly similar localization landscapes, which also resemble DTT-treated ATAC-Seq profiles.

E) Correlation matrix between CUT&Tag and ATAC libraries. Datasets for untreated sperm are all distinct, whereas ATAC-Seq and CUT&Tag datasets for permeabilized sperm are broadly concordant, consistent with CUT&Tag data simply reflecting untargeted Tn5 activity in the CUT&Tag protocol.

**Supplemental Figure S5. Features of sperm ChIP-Seq datasets**

A) Heatmaps of H3K4me3 and H3K27me3 bivalent data for the DTT only and DNase+DTT libraries, aligned over all promoter regions (n = 57,102;TSS +/− 2.5 kb), organized from high to low enrichment of H3K4me3 signals.

B) Sequence logos for enriched motifs at CTCF ChIP-Seq peaks called from DTT-only and DNase+DTT libraries, as indicated.

C) Limited solubilization of the sperm genome following sonication. Formaldehyde-fixed sperm were sonicated to fragment the genome for ChIP-Seq. Following sonication, resulting material was centrifuged (10,000 xg, 10 min) and DNA was extracted from pellet and supernatant fractions and characterized by gel electrophoresis.

D) Native MNase-ChIP-Seq for H3K4 and K27 methylation. Data for untreated sperm closely match previously-reported histone modification profiles; curiously, for this assay – unlike for other histone modification mapping protocols – we find essentially no change following DNase or DTT treatments. We speculate that DNase treatment of cell-free chromatin leaves some nucleosomes intact, which are available for immunoprecipitation but which do not have adjacent linker DNA required for Tn5 insertion in CUT&Tag, or which add to a global nonspecific background for CUT&RUN where MNase-digested nucleosomes are used for readout.

**Supplemental Figure S6. Hi-C scaling plots**

Top panel shows interaction frequency between two loci at increasing genomic distances (x axis) for the indicated datasets. Bottom panel shows the derivative of these curves. The relatively flat profile seen in the DNase+DTT dataset is typical for poorly-crosslinked Hi-C libraries, potentially owing to the paucity of lysines in sperm DNA-associated proteins like the protamines.

We note that although the majority of published Hi-C studies of mouse sperm exhibit typical somatic features including A/B compartments and TADs (Battulin et al. 2015; Jung et al. 2017; Ke et al. 2017; Alavattam et al. 2019; Jung et al. 2019; Wang et al. 2019b), TADs were notably absent in (Vara et al. 2019). Although this would be explained parsimoniously had DTT been used in that study – generating maps dominated by bona fide sperm rather than contaminating chromatin, as seen in our **Fig. 6** data – this does not appear to have been the case based on the Vara *et al* Methods section.

Instead, two technical choices may provide a potential explanation. First, Vara *et al* state that testis, epididymis, and epididymal sperm were co-incubated during testicular dissociation, conditions which include DNase I treatment (DNase treatment is a typical part of testicular dissociation protocols). Of course this would eliminate the cell-free chromatin contamination present in the cauda epididymis, as demonstrated throughout this manuscript, but raises the question of what material was captured in the resulting Hi-C libraries. This highlights the second technical choice, where Vara *et al* snap freeze FACS-sorted sperm prior to all fixation and permeabilization steps. In our experience, MNase-Seq data are dramatically altered when the assay is performed on frozen and thawed sperm vs. freshly isolated sperm used immediately (not shown). We speculate that this reflects ice damage to sperm frozen without cryoprotectant, potentially allowing later enzyme access to fractured sperm heads. In this scenario Vara *et al* would be able to generate Hi-C maps, free of cell free chromatin, from sperm broken by freezing.

