## Supplementary figures and images for "Revisiting chromatin packaging in mouse sperm"

### Fig S1

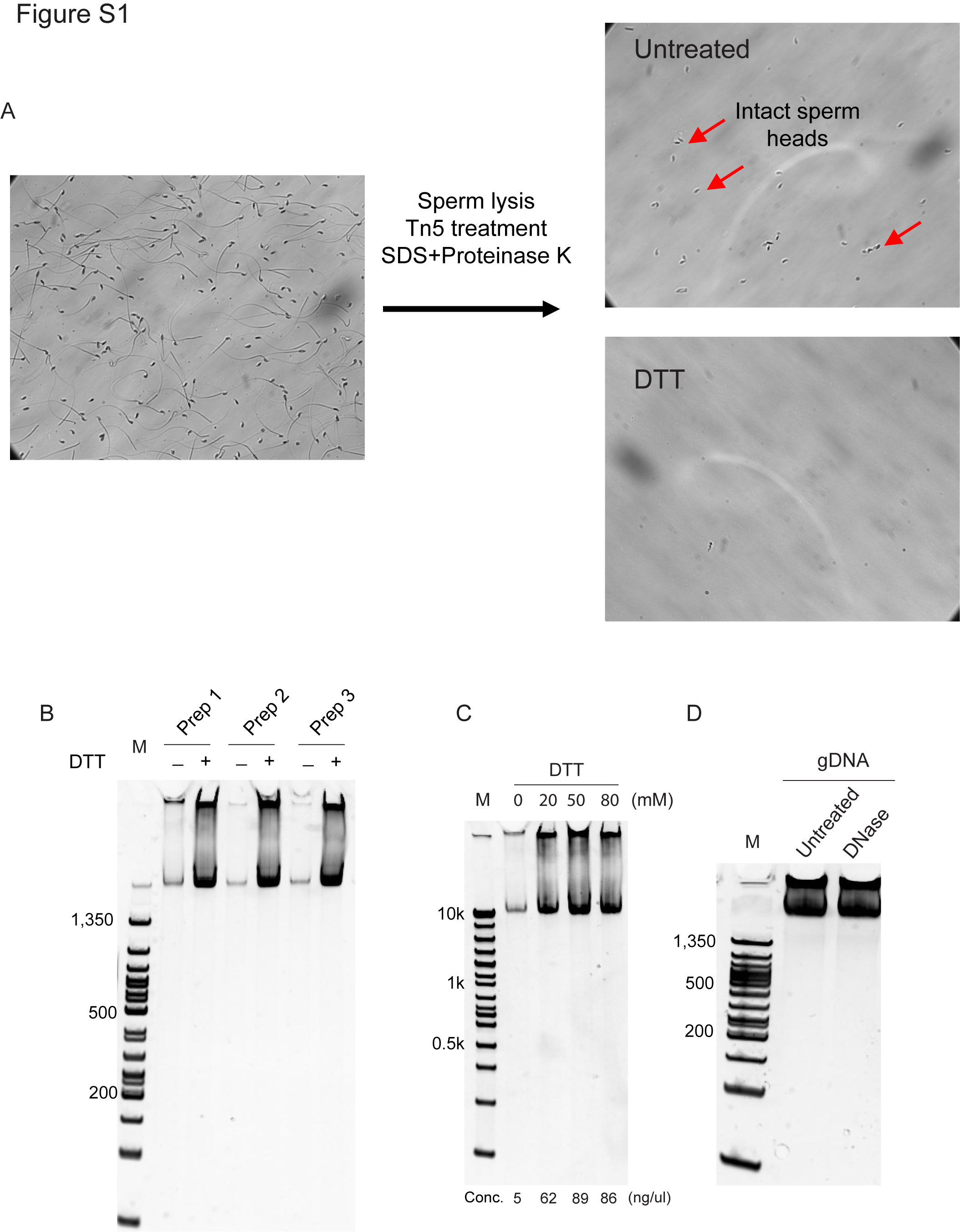

### Fig S2

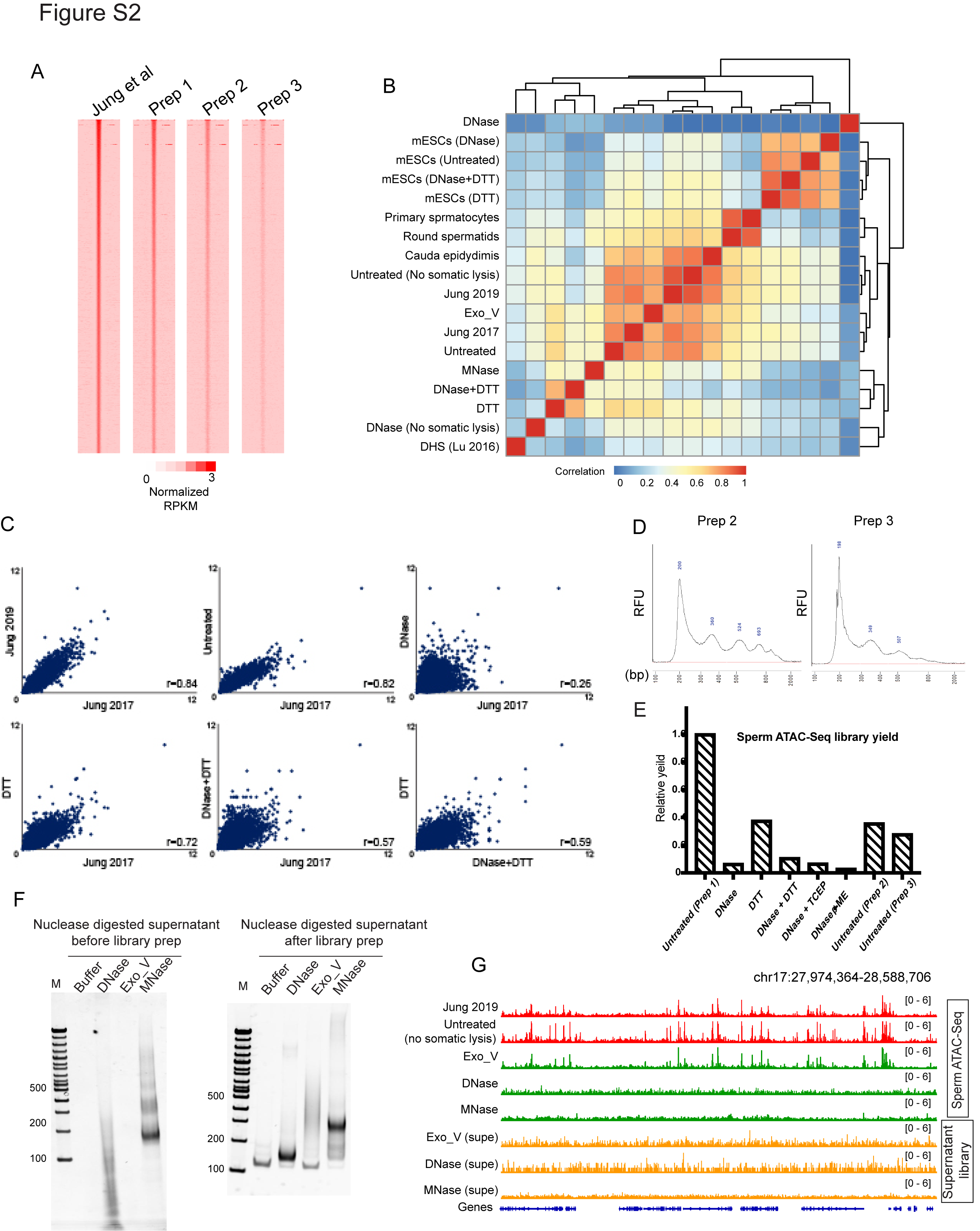

### Fig S3

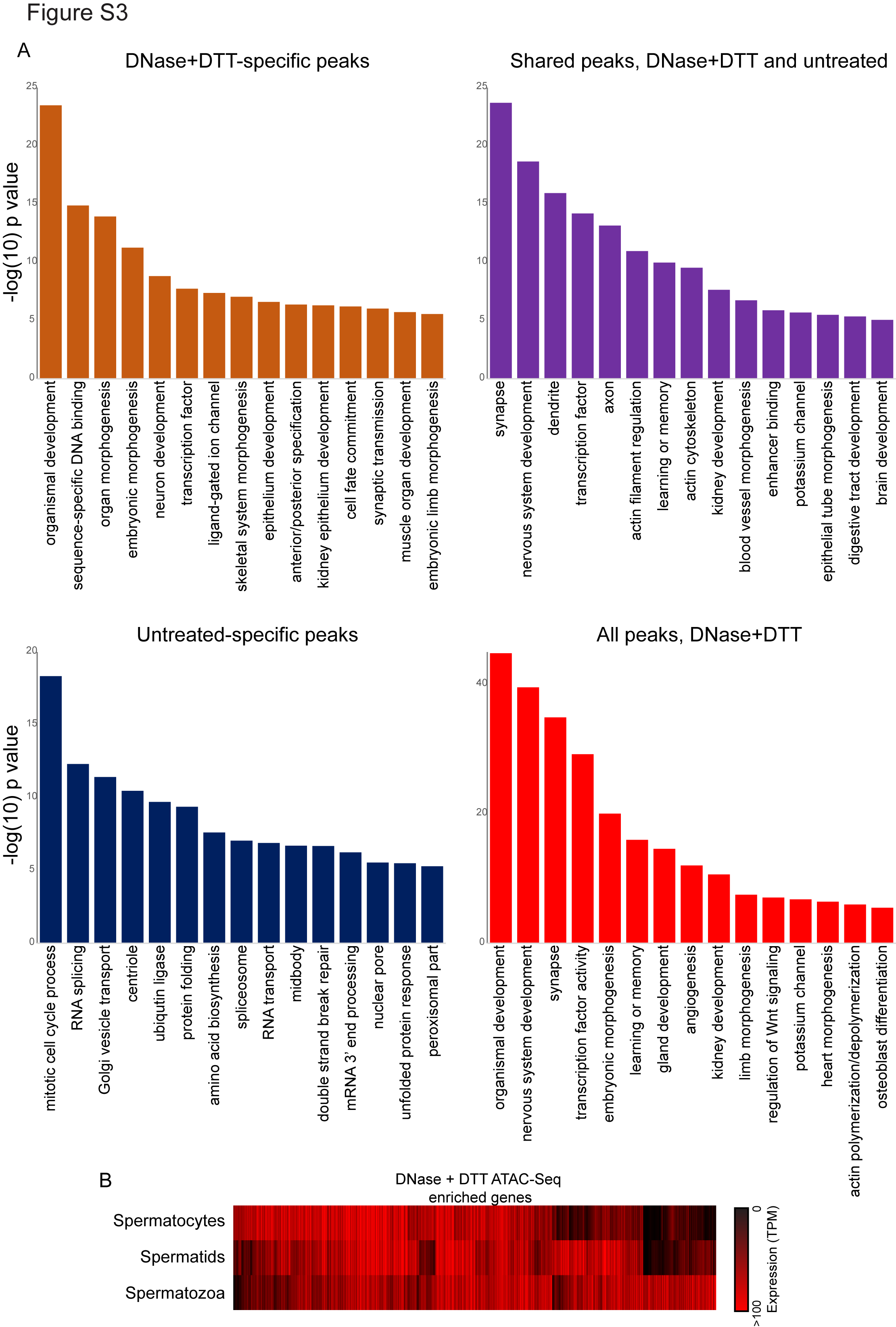

### Fig S4

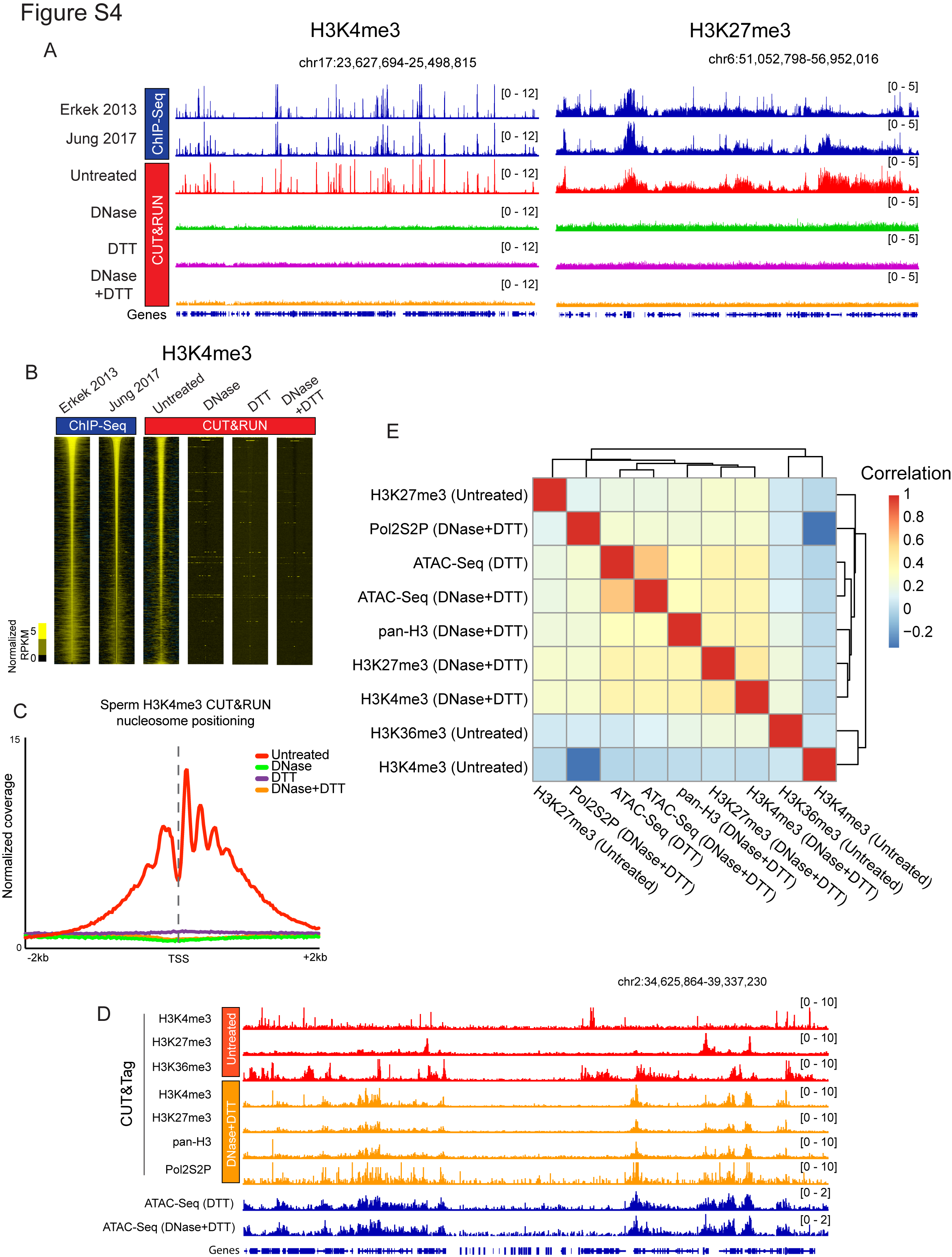

### Fig S5

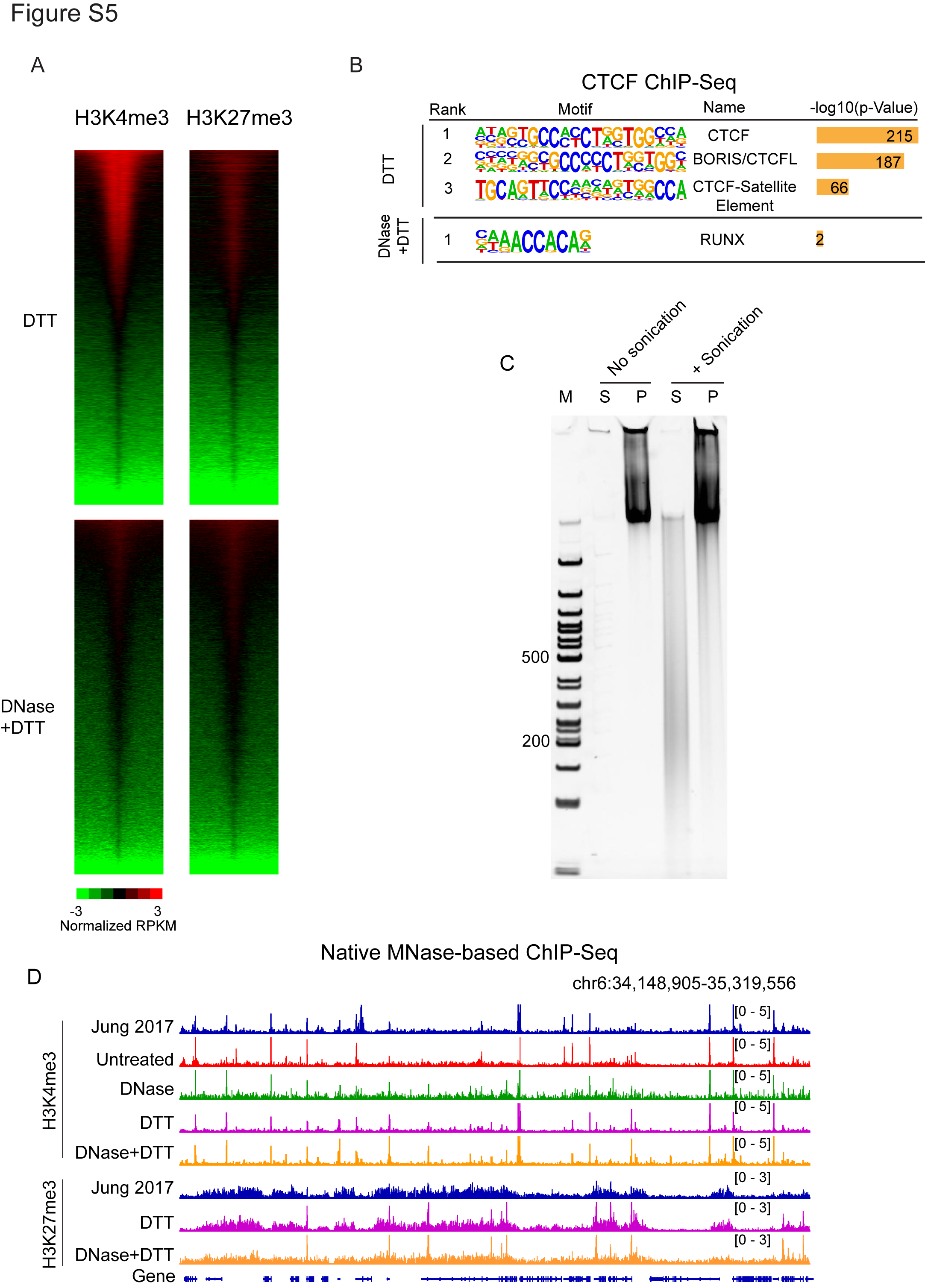

### Fig S6

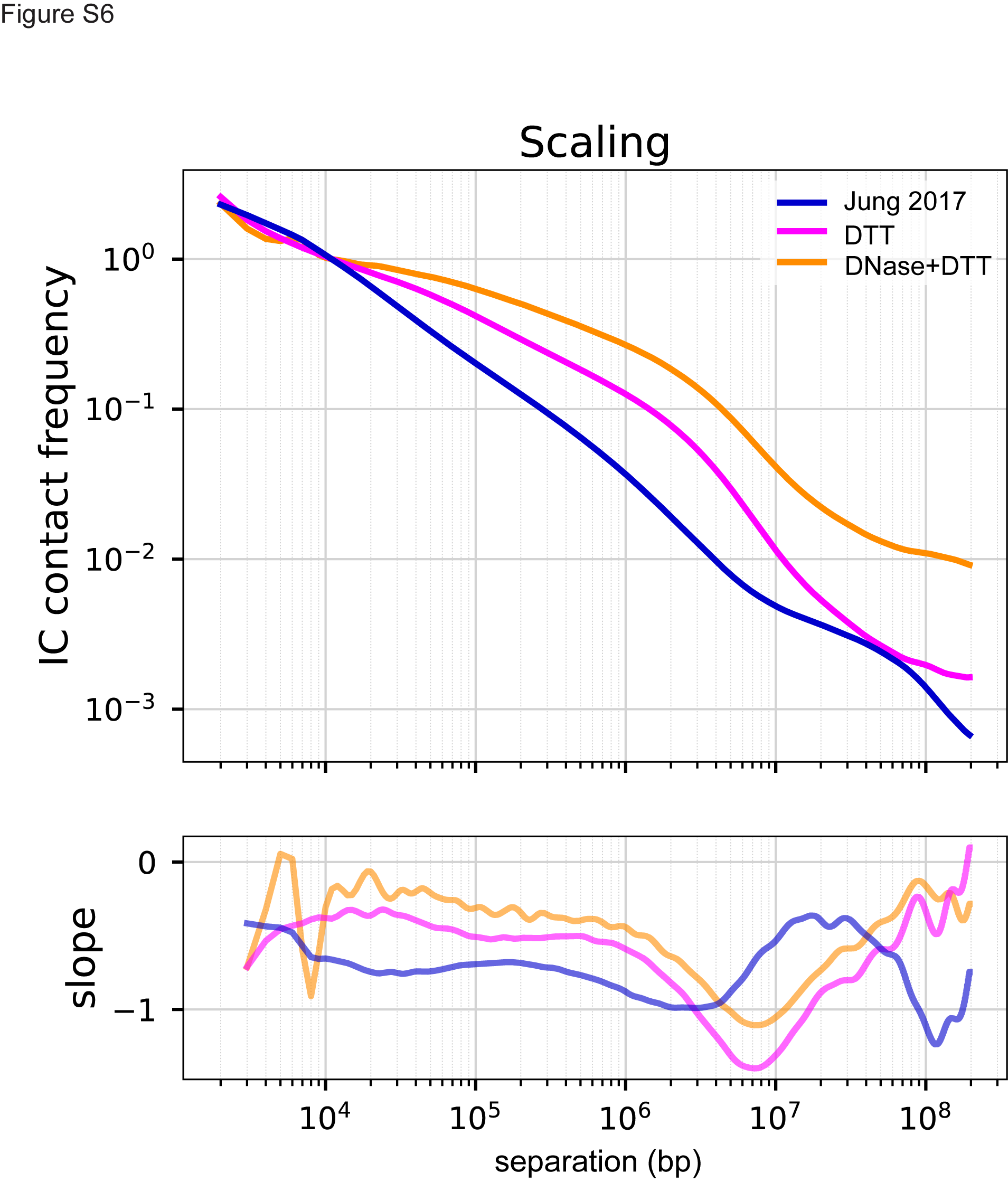
